# Cell-Derived Vesicles as TRPC1 Channel Delivery Systems for the Recovery of Cellular Respiratory and Proliferative Capacities

**DOI:** 10.1101/2020.05.16.099283

**Authors:** Felix Kurth, Yee Kit Tai, Dinesh Parate, Marc van Oostrum, Yannick R. F. Schmid, Shi Jie Toh, Jasmine Lye Yee Yap, Bernd Wollscheid, Alaa Othman, Petra S. Dittrich, Alfredo Franco-Obregón

## Abstract

Pulsed electromagnetic fields (PEMFs) are capable of specifically activating a TRPC1-mitochondrial axis underlying cell expansion and mitohormetic survival adaptations. This study characterizes cell-derived vesicles (CDVs) generated from C2C12 murine myoblasts and shows that they are equipped with the sufficient molecular machinery to confer mitochondrial respiratory capacity and associated proliferative responses upon their fusion with recipient cells. CDVs derived from wild type C2C12 myoblasts include the cation-permeable transient receptor potential (TRP) channels, TRPC1 and TRPA1, and directly respond to PEMF exposure with TRPC1-mediated calcium entry. By contrast, CDVs derived from C2C12 muscle cells in which TRPC1 had been genetically knocked-down using CRISPR/Cas9 genome editing, do not. Wild type C2C12-derived CDVs are also capable of restoring PEMF-induced proliferative and mitochondrial activation in two C2C12-derived TRPC1 knockdown clonal cell lines in accordance to their endogenous degree of TRPC1 suppression. C2C12 wild type CDVs respond to menthol with calcium entry and accumulation, likewise verifying TRPA1 functional gating and further corroborating compartmental integrity. Proteomic and lipidomic analyses confirm the surface membrane origin of the CDVs providing an initial indication of the minimal cellular machinery required to recover mitochondrial function. CDVs hence possess the potential of restoring respiratory and proliferative capacities to senescent cells and tissues.

## 1. Introduction

Cellular aging is regulated by mitochondrial respiratory capacity.^[1]^ Conversely, cellular senescence arises from mitochondrial functional deterioration, a process known as mitochondrial dysfunction-associated senescence (MiDAS).^[2–4]^ We have previously shown that mitochondria are capable of responding to environmental stimuli by their communication with cell surface-associated signaling complexes and, via such an arrangement, cellular aging can be influenced by externally arising stimuli such as mechanical forces, temperature, or magnetic fields.^[5]^ The use of cell-derived vesicles (CDVs) as delivery vehicles for cellular sensory apparatuses is thus appealing as they have been shown in some cases to retain the necessary molecular components to form functioning signaling apparatuses capable of conferring cellular responses to environmental stimuli.^[6–9]^ CDV-based platforms may hence be applicable to reverse states of cell senescence or quiescence as a result of mitochondrial dysfunction.^[2,3]^ This objective had only been previously approached with the use of native extracellular vesicles (exosomes and microvesicles) produced by mesenchymal stem cells, but with limited success largely due to endogenously low yields of biological material,^[10–13]^ a limitation that would not prove as restrictive to CDV production. Moreover, CDV-based platforms are not subject to certain caveats associated with conventional cell-based assays such as nutrient requirement, regulated viability or dynamic changes in signaling capacity due to cell cycle progression, differentiation, or senescence.^[5,14]^ Instead, CDVs would represent a snapshot of the cells’ physiological state at the time of vesicle formation.

Different protocols of producing CDVs have been previously developed. Sufficiently strong shear forces are capable of producing membrane vesicles, yet of relatively poorly defined composition due to the non-specific nature of the shedding forces.^[15–17]^ Inducing membrane blebbing with the use of cytochalasin B treatment alleviated the need for harsh mechanical disruption as well as improved CDV yield and functionality by allowing plasma-membrane signaling microdomains to remain intact.^[8,9,18]^ Nonetheless, although CDVs produced in this manner are generally assumed to retain native plasma-membrane structure and integrity, detailed information of the proteome and liposome composition of CDVs remains sparse. Here, we provide an initial in-depth characterization of CDVs derived from murine muscle cells produced via a variation of the cytochalasin B method in terms of yield, size, lamellarity as well as protein and lipid compositional analyses. We also demonstrate the presence of molecular signaling apparatuses capable of responding to external stimuli as well as conferring functionality to naïve cells.

Cellular sensory perception is initiated at the cell periphery and is enabled by surface membrane-embedded molecular signal transduction apparatuses. Transient receptor potential (TRP) channels have been implicated in diverse modes of cellular sensory transduction including temperature, light, redox, pH, mechanosensation, and magnetoreception.^[5,19,20]^ TRPC1 and TRPM7 are the most abundant and ubiquitously expressed of all TRP channels and of clear developmental importance.^[21]^ In particular, TRPC1 expression has been implicated in calcium-dependent proliferative responses of skeletal muscle progenitor cells to mechanical or magnetic stimulation.^[5,22–25]^ Notably, magnetically-induced TRPC1-mediated calcium entry has been recently shown to activate mitochondrial respiration and downstream cellular survival adaptations via a process of magnetic mitohormesis.^[5]^ TRP channel activation has been traditionally studied in intact cell systems where the minimum molecular requirements for the channel to fulfill its endogenous cellular role has been problematic to decipher. Therefore, the nuances distinguishing channel gating and channel-assisted cellular responses have been difficult to adequately address. Given the fact that TRP channel responses have also been observed in planar lipid bilayers, however, indicates that some TRP channel isoforms are capable of independent gating.^[26]^ Nonetheless, planar lipid bilayers do not recapitulate the mechanical nor molecular complexity of the native plasma membrane that may be necessary for functional biological sensor development nor would they be capable of acting as fusogenic delivery-vehicles for ultimate use in cell therapies.^[27]^ A major objective of this study was hence to test for the presence and functionality of TRP channels within isolated native skeletal muscle-derived CDVs and, if so, to ascertain whether CDVs demonstrate the sufficient molecular machinery to recover loss of cellular sensory perception and mitochondrial respiration in deficient cells as an initial indication of minimal molecular sensory apparatus requirement to fuel future studies **(Figure 1)**.

**Figure 1.**
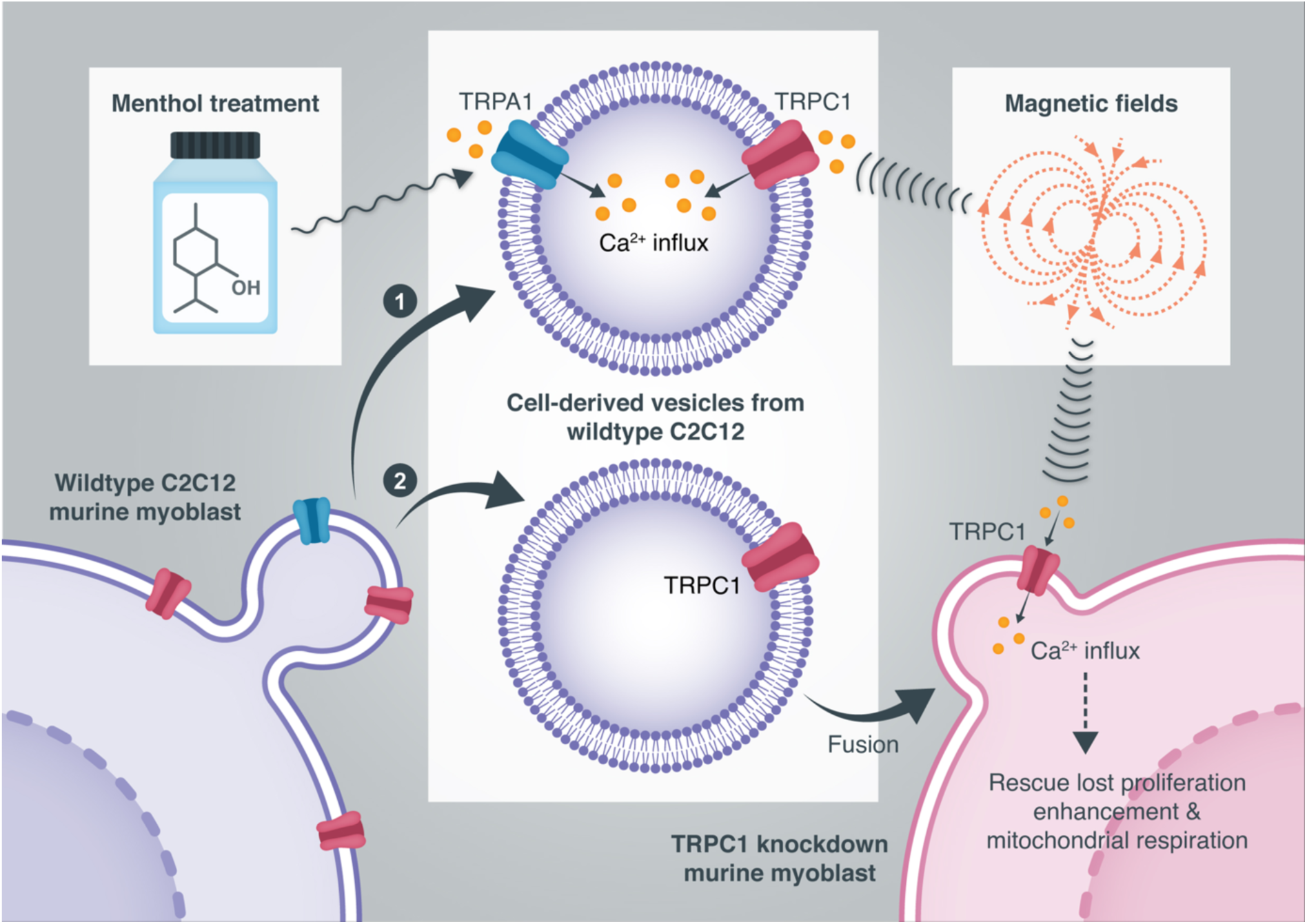
Schematic depicting the tests applied to CDVs to ascertain (**1**) innate channel functionality and (**2**) recuperative potential for cellular respiratory and proliferative capacities. **1**) Cell-free assays were comprised of examining specific TRP channel activation by menthol (TRPA1) or magnetic fields (TRPC1). **2**) Cell-based assays entailed administering wild type CDVs to TRPC1-knockdown myoblasts followed by testing for recuperation of cellular proliferative and mitochondrial respiratory capacities.

## 2. Results

### 2.1. Characterization of liposome formation

The cytochalasin B method of producing CDVs is an established protocol.^[6–9]^ Nonetheless, vesicle yields employing this method vary with cell type, culture conditions or subtle variations in the preparation protocol.^[8,9]^ Dynamic light scattering (DLS) measurements indicated a ∼10-fold greater yield of CDVs with the addition of a shaking step (300 rpm, 37 °C, 15 min) implemented during the incubation with cytochalasin B. **Figure 2A** (left) depicts CDV yields as a function of cell density and myogenic differentiation status. CDV count scaled up linearly with increasing density of individual myoblasts as well as following their fusion into differentiated syncytial myotubes. On the other hand, lowering the osmolarity (∼33%) of the incubation and hosting solutions reduced the yield of CDVs, particularly under static conditions (∼3-fold; data not shown). Although the size distributions of CDVs derived from either myoblasts or myotubes were not statistically different (Figure 2A, right), CDVs from myotubes exhibited a trend to be larger (Figure 2B, left). Extrusion narrowed the size distribution of CDVs;^[28,29]^ representative size distributions with and without extrusion are shown in Figure 2B (right). Once extruded, CDVs remained stable in size for several weeks.^[28]^

**Figure 2.**
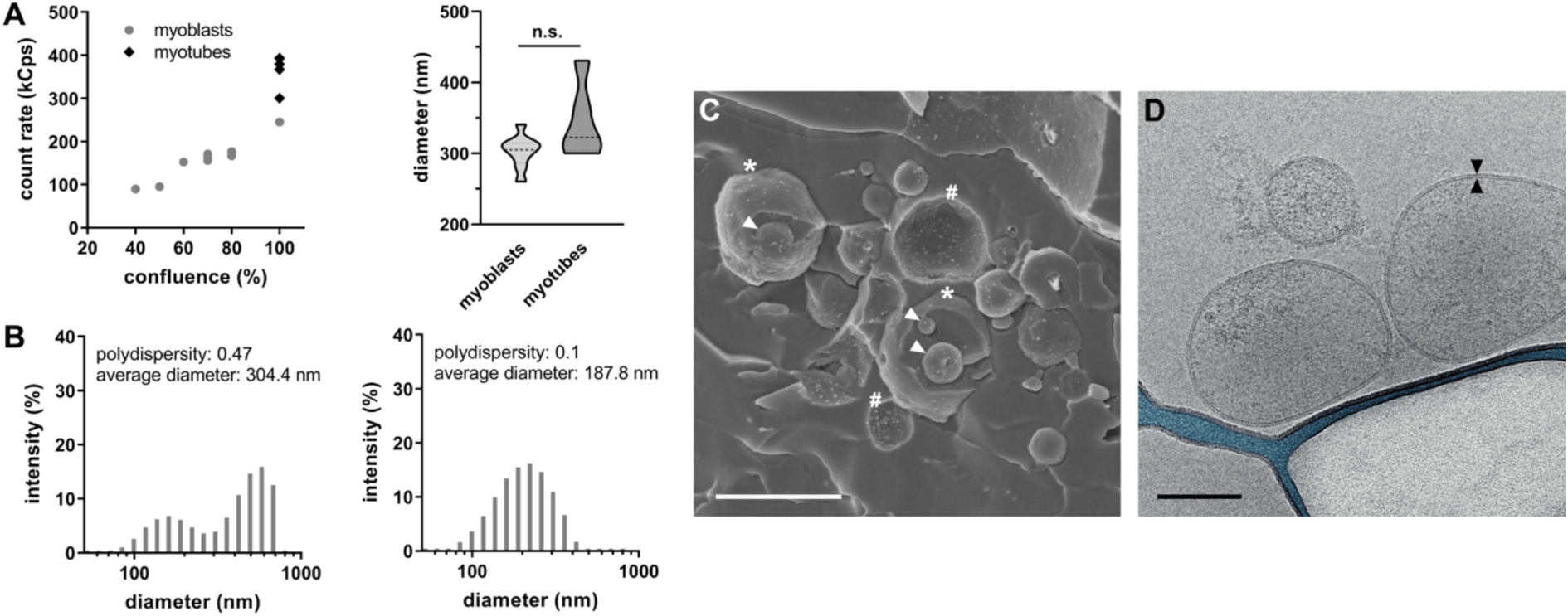
(A) CDV yield is influenced by cell confluence and cellular differentiation status. Vesicle yield increased with increasing confluence and in differentiated myotubes (left). Mean CDV diameter was not statistically different in samples generated from myoblasts or myotubes (right) (n.s.: not significant). (B) CDV size distribution without (left) and with extrusion (right) through 200 nm diameter pores. (C) Freeze fracture SEM of a non-extruded CDV sample. Asterisks (*) identify liposomes that partially look out of the surface, hashtags (#) depict imprints within the surface from CDVs that were entirely detached during the fracture process thereby leaving a crater in the surface. Arrows reveal smaller liposomes that were enclosed within the lumens of larger liposomes during the CDV formation. Scale bar: 1 µm. (D) TEM micrograph of extruded CDVs shown in B (right). The blue colored area depicts the free carbon grid. The black triangles mark the lipid bilayer with a diameter of 4 nm to 5 nm; individual monolayers are visible as thin dark lines. The 2 CDVs adjacent to the free carbon grid are deformed due to the limited height of the sample holder. Scale bar: 100 nm.

Electron microscopic analysis of non-extruded CDVs revealed diameters ranging between a few tenths of nanometers to >1 µm, coinciding with DLS measurements. Larger, non-extruded vesicles were often found to enclose smaller vesicles, possibly exosomes, that may have originated from components of the endosomal or reticular compartments proximal to the surface membrane that was captured at the time of vesicle formation (Figure 2C). Smaller extruded CDVs (∼200 nm) characteristically exhibited unilamellarity by cryo-transmission electron microscopy (Figure 2D) and internal proteinaceous granularity, *cf.* ^[30]^, previously suggested to pertain to captured cytoplasm (also see Figure S1B and C). The cytochalasin B method thus produces CDVs that may contain remnants of membrane-associated intracellular scaffolded signaling-complexes.

### 2.2. Proteome analysis

CDV samples were subjected to mass spectrometry-based proteomic analysis that served to identify a total of ∼1200 protein groups (Supplementary file Proteotype_CDV_C2C12.xlsx). Comparing the CDV proteotype data to the deepest available proteome data base for C2C12 myoblasts ^[31]^ and segregating them according to cellular compartments using Gene Ontology (GO) revealed that CDVs were relatively enriched in proteins associated with the plasma membrane and extracellular domains, whereas protein species characteristic of intracellular compartments were underrepresented (**Figure 3A**). Succinctly, CDV-derived proteins were enriched in molecular moieties associated with cellular stimuli-reception and transduction, whereby proteins associated with peripheral cellular compartments (plasma membrane and extracellular regions) represented ∼32% of all identified proteins (Figure 3B), ∼ 2-fold greater compared to intact C2C12 cells. More detailed summaries of the assigned molecular functions, biological processes as well as GO-cell analyses are given in Figure S2 and Table S1, respectively.

**Figure 3.**
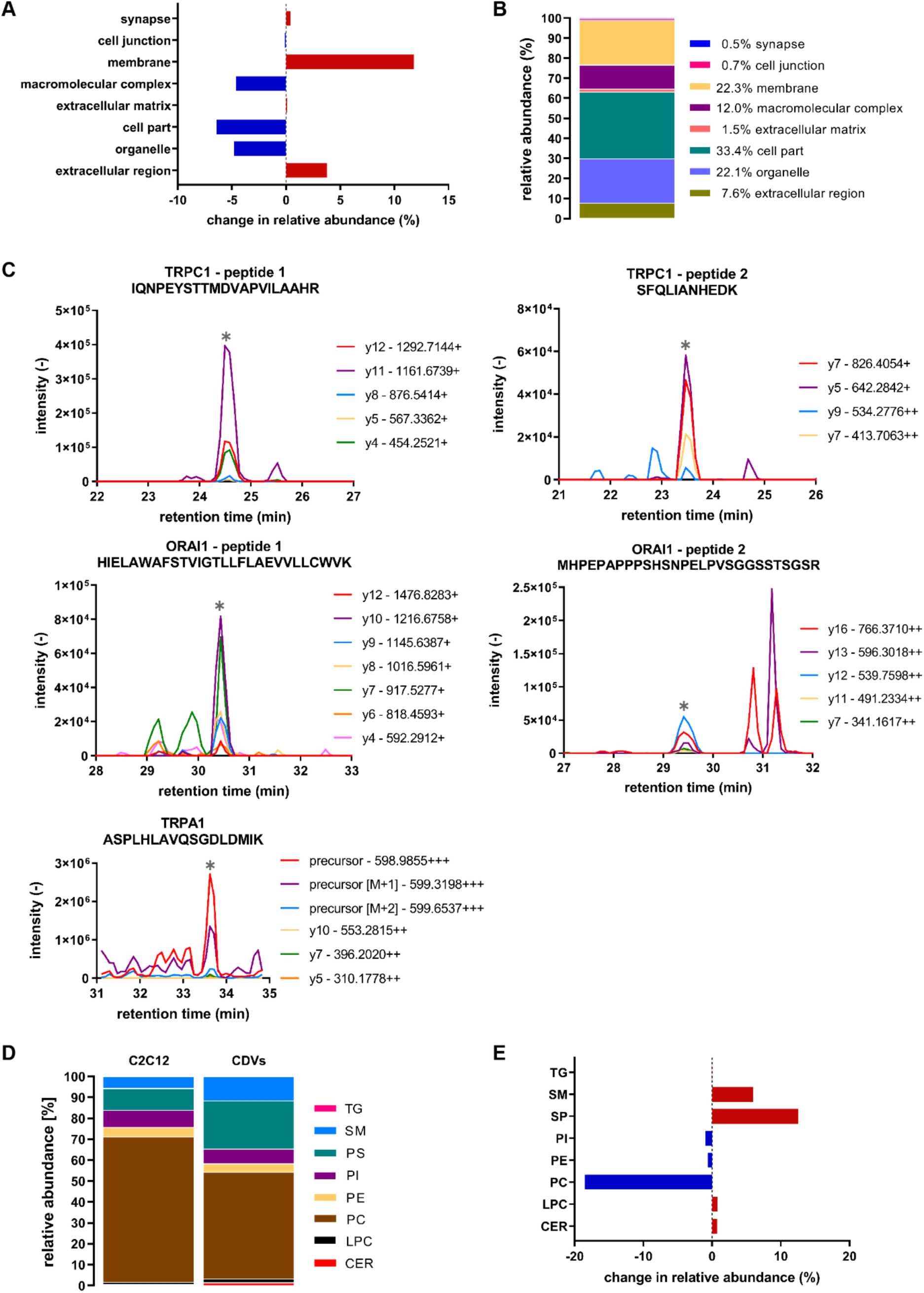
(A) Proteome analysis of CDVs and its comparison to published data.^[31]^ Differences in GO-assigned fractions of identified proteins in between CDVs and C2C12 derived from myotubes. CDVs exhibit relative enrichments of protein identifications with GO terms associated with membrane and the extracellular region. (B) The percentage of total identified proteins for GOs of CDV samples provides an overview of the cell component protein associations. (C) Chromatographic traces (PRM measurements) of fragment ions matching proteotypic peptides from the respective calcium channels TRPC1, ORAI1, and TRPA1. Grey asterisks mark the relevant signals. (D) Lipid abundances for CDVs and C2C12 myotubes. (E) Comparison of lipid abundances between CDVs and C2C12 myotubes. The most significant changes in lipid content (CDVs relative myotubes) originates from the sphingomyelins (+5.99%) and phosphatidylserines (+12.5%) as well as from phosphatidylcholines (−18.51%). A list of the detailed concentrations of lipid components in CDVs and whole cell samples is available in the appendix. Lipid abbreviations are: TG: triglyceride, SM: sphingomyelin, PS: phosphatidylserine, PI: phosphatidylinositol, PE: phosphatidylethanolamine, PC: phosphatidylcholine, LPC: lysophosphatidylcholine, CER: ceramide.

TRPC1 stimulates mitochondrially-stimulated myogenesis,^[5]^ downstream of TRPC1-mediated calcium entry.^[22,23,32,33]^ Intramyocellular calcium levels are adjusted by a delicate interplay between plasmalemmal calcium influx channels, such as TRP channels, and intracellular calcium store release channels that are mutually modulated via a process of Calcium-Induced Calcium Release (CICR).^[33–35]^ Foremost, TRPC1 channels in combination with ORAI1 have been implicated in CICR.^[22,34–39]^ TRPA1 has also been described to contribute to CICR.^[40]^ In order to ascertain TRP channel expression in CDVs, parallel reaction monitoring (PRM) was used to specifically detect proteotypic peptides mapping to TRPC1, ORAI, TRPA1, TRPV1 or TRPV2. Co-eluting peaks of peptide fragments matching those of proteotypic peptides originating from TRPC1, ORAI1, and TRPA1 were identified (Figure 3C). By contrast, we could not detect matching proteotypic peptides for TRPV1 and TRPV2 previously shown to be transcriptionally represented in C2C12 myoblasts.^[5,23]^ TRPA1 activity can be induced by cold temperature,^[41,42]^ whereas TRPC1 can be activated by either mechanical forces ^[5,22]^ or by low amplitude magnetic fields.^[5]^ Magnetic field activation of TRPC1 will be employed here for their direct (diffusion-independent), non-contact, and immediate mode of delivery.

### 2.3. Lipid analysis

The surface membrane of intact cells is enriched in the sphingomyelins (SM), phosphatidylserine (PS) and cholesterol, whereas phosphatidylcholine (PC) is expressed in relatively lower abundance.^[43]^ On the other hand, phosphatidylethanolamine (PE) and phosphatidylinositol (PI) do not differ significantly in relative abundance between the distinct membrane compartments. Lipid analysis corroborated the surface membrane origin of our CDVs (Figure 3D, E), by demonstrating relative enrichments in SM and PS and depletion of PC.^[9,18]^ Notably, products of the sphingomyelin pathway have been implicated in modulating TRPC1 function upstream of myogenic progression ^[44,45]^ and are a major constituent of surface lipid rafts,^[46]^ wherein TRPC1 assumes its sensory transduction role upstream of CICR,^[37– 39,47,48]^ providing relevance to their relative enrichment in CDVs. Moreover, as fusogenic capacity is conferred upon differentiating myoblasts by uniquely elevated surface membrane expression of PS,^[49]^ fusogenicity could be potentially transferred to isolated CDVs and be a specific functionality of myoblast-derived CDVs. A detailed breakdown of CDV lipid composition is given in Table S2.

### 2.4. CDV functionality

To ascertain TRP channel functionality in our CDVs, we characterized their thermal and magnetic responses. We first established CDV integrity by employing a fluorescent calcein dye covalently functionalized with the addition of an acetoxymethyl (AM) group, conferring the dye with lipid solubility. Upon partitioning into an intact and viable cellular membrane compartment, however, enclosed endogenous esterases enzymatically cleave the calcein-AM ester linkage, entrapping the calcein therein for subsequent detection. CDVs, immobilized onto the substrate of a microfluidic channel via printed cholesterol linkers ^[50]^ (**Figure 4A, 1**), were loaded with calcein-AM and imaged by fluorescence microscopy (Figure 4A, 2). Figure 4B shows microcontact-printed cholesterol-linkers that had been conjugated onto the substrate of a microfluidic channel and visualized with Texas Red-avidin (red; left). Cholesterol-immobilized CDVs exhibited fluorescence, indicating that the dye had been successfully internalized, enzymatically cleaved and captured for microscopic imaging (green, right). TRPA1 channel functionality was next ascertained by exploiting its inherent sensitivity to menthol, a TRPA1 channel agonist that emulates cold temperature at the level of channel transduction.^[41,42,51]^ Immobilized CDVs were loaded with the calcium-sensitive dye, Fluo-4 AM and challenged with menthol (500 µmol L^-1^), inducing a robust increase in Fluo-4 fluorescence that reflected menthol-stimulated, TRPA1-mediated, calcium uptake into CDVs (Figure 4C). This result would also indicate that at a majority of the CDVs exhibited sufficient compartmental integrity to restrict the escape of dye and calcium through the duration of the experiment.

**Figure 4.**
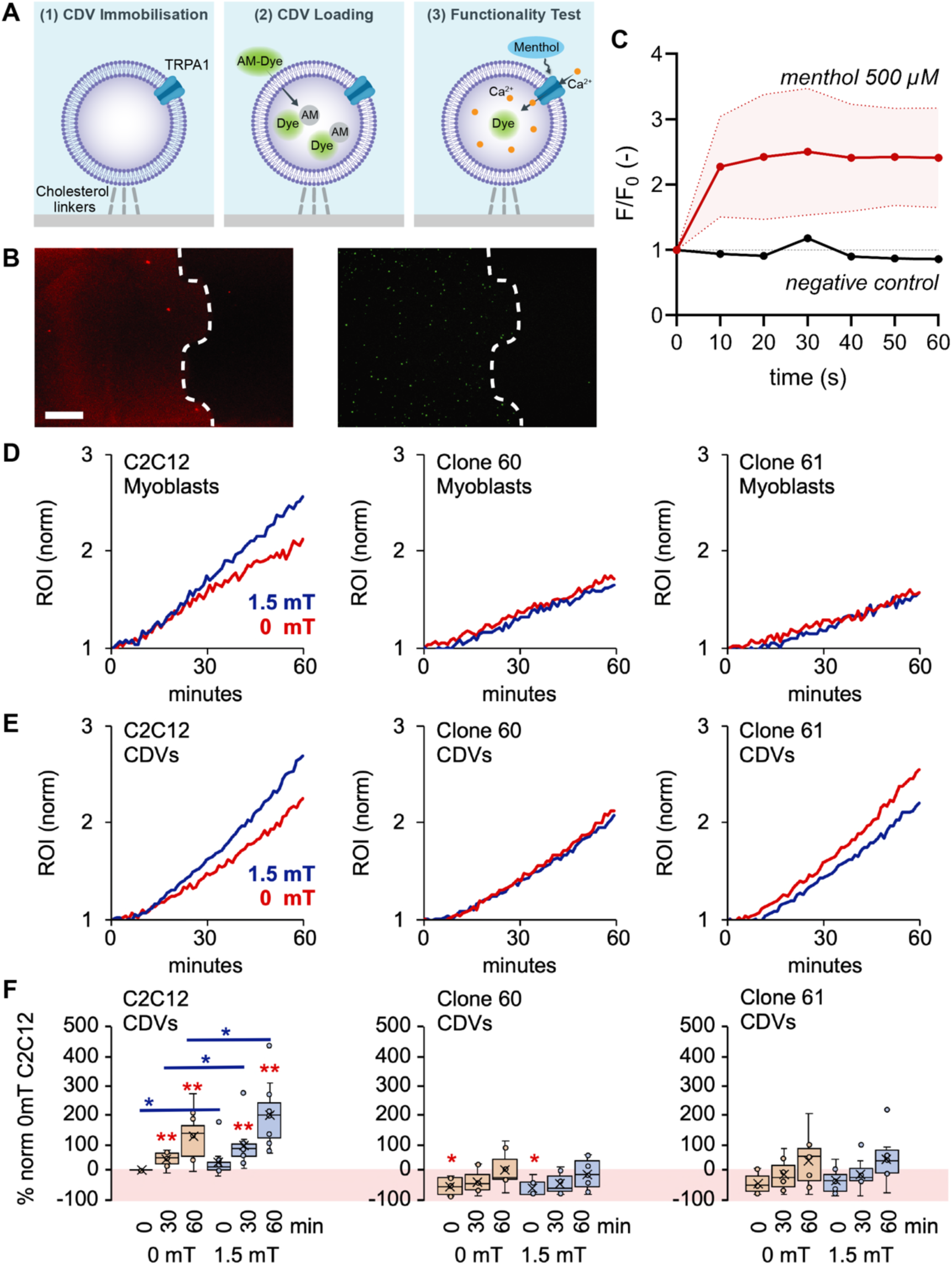
(A) Schematics of the CDV immobilization (1), loading (2) and TRP channel functional assessment (3) (B) Micrographs of the print of the biochemical cholesterol linkers (left) and CDVs immobilized on the same print and loaded with calcein-AM. Red fluorescence is Texas Red as part of the biochemical linker, the dotted line indicates the edge of the print. Scale bar: 20 µm. (C) Examples of Menthol-induced increase in intra-CDV calcium concentration (measurement in 2 independent microfluidic channels, red data trace) compared to a control condition without menthol administration (negative control; black data trace). Data was recorded simultaneously in individual, parallel channels on a microfluidic chip using CDVs from the same batch. The negative control signal represents the average value within a single microfluidic channel, the menthol signal represents the average ± standard deviation (shaded area) of 2 individual microfluidic channels. (D-E) TRPC1-mediated calcium uptake in response to magnetic stimulation. Representative calcium responses (Rate of Rise (ROR) normalized to starting value) from intact C2C12 myoblasts (D) or CDVs (E) derived from wild type C2C12 myoblasts (left) or C2C12 CRISPR/Cas9 knockdown clones, 60 (middle) and 61 (right), in response to brief (10 min) magnetic field exposure (1.5 mT) while in suspension immediately before reading. Blue and red lines represent magnetically-exposed and control samples, respectively, as indicated. (F) Mean calcium responses from CDVs derived from C2C12 wild types (left) or knockdown clones 60 (middle) and 61 (right) as indicated. All values are shown normalized to C2C12 CDVs at time 0 min and 0 mT. CDVs were loaded with Calcium Green-1 AM (0.5 ng/µl) for 30 min while in culture followed by preparation of CDVs for subsequent magnetic stimulation and immediate analysis. Red shaded area reflects inhibition relative to C2C12 CDVs at time 0 min and 0 mT. Data represents the means of 6-11 independent experiments (biological replicates), each representing the means of 8-10 technical replicates. ** and * represent *P* < 0.01 and < 0.05 relative to respective 0 mT at same time point (blue) or to time = 0 min in C2C12 CDVs (red) as indicated. Statistical significance was not achieved in either TRPC1-knockdown clone, 60 or 61, relative to their respective 0 mT or 0 min conditions.

TRPC1-mediated calcium entry has been shown to be specifically induced by brief exposure to non-invasive pulsing magnetic fields (PEMFs), upstream of mitochondrial activation in muscle cells.^[5]^ This effect could be elicited from myoblasts in suspension and therefore, independent of other sources of cell-to-cell or cell-to-substrate activation, allowing for a more precise comparison of responses from intact cells and membrane-derived CDVs. C2C12 myoblasts cultures were loaded with the calcium-sensing dye, Calcium Green-1 AM, followed by enzymatic detachment from the substrate and brief (10 min) exposure to PEMFs while in suspension. Intact wild type C2C12 myoblasts responded to magnetic stimulation with calcium entry, *cf.* ^[5]^ (Figure 4D, left), whereas genetically-engineered C2C12 myoblast clonal cell lines (c60 and c61) in which TRPC1 expression had been knocked down using CRISPR/Cas9 genetic silencing ^[5]^ were incapable of responding in a like manner to similar magnetic field stimulation (Figure 4D, middle and right). We next tested whether magnetoreception could be isolated to membrane-derived CDVs and therefore, a property of the surface membrane, rather than an integrated response of the intact cell system. Only CDVs derived from wild type C2C12 muscle cells (Figure 4E, left), but not identically-loaded TRPC1 knockdown clones, c60 and c61 (Figure 4E, middle and right, respectively), were capable of responding to magnetic stimulation with increased calcium fluorescence, effectively recapitulating the intact cell response. Additionally, basal (0 mT) calcium levels were also lower in CDVs derived from TRPC1 knockdown clones relative to wild type CDVs (Figure 4F), validating previous assertions that TRPC1 channels are implicated in establishing cellular basal calcium levels.^[5,22]^ These findings corroborate the presence of TRP channels in CDVs as well as demonstrate that TRPC1 per se is sufficient to confer magnetoresponsive functionality for potential delivery and recovery of magnetoreception in recipient cells deficient in said processes or for the development of biosensing platforms aimed at detecting magnetic fields.

### 2.5. Liposome-cell fusion

Myoblasts are a fusogenic cell type due to inherently elevated PS expression levels in their outer membranes,^[49]^ an attribute that can be potentially translated to our specific CDV preparation due to their muscle origin and relative membrane enrichment (Figure 3D). CDV-cell fusion-mediated membrane incorporation was examined to ascertain the potential for the functional transfer of signaling complexes to recipient cells. CDVs were labeled with the lipophilic membrane dye R18 (1 µmol L^-1^) and then added to C2C12 myoblasts cultures. Thirty minutes after administration of CDVs to myoblast cultures, intracellular staining arising from labeled membranes could be detected by confocal microscopy, particularly to perinuclear regions (Figure S3). Although, TRPC1 function has also been ascribed to intracellular localizations in skeletal muscle cells,^[52]^ it is unknown whether here TRPC1 colocalized with intracellular membrane staining or intracellular membrane recycling was occurring in isolation. Nonetheless, our reported uptake and intracellular transportation times coincide with those previously reported for exosomes, including membrane trafficking to perinuclear localizations.^[53]^ The macromolecular organization and operation of the membrane-associated fusogenic apparatus ^[27]^ as well as the cellular physiological consequences of CDV internalization will be the focus of future investigations.

### 2.6. Functional rescue of myoblast proliferative capacity

We next tested whether CDVs could functionally rescue loss of magnetically-sensitive proliferation as a result of genetically targeted knockdown of TRPC1. Cultures of wild type and TRPC1 knockdown C2C12 myoblasts were exposed to PEMFs for 10 min and allowed to proliferate for 24 h before growth assessment **(Figure 5A)**. Under control conditions (−CDVs) magnetic stimulation enhanced the proliferation of wild type C2C12 myoblasts (Figure 5A, top), whereas the TRPC1 knockdown myoblast clones, c60 (Figure 5A, middle) and c61 (Figure 5A, bottom), instead exhibited depressed proliferation in response to magnetic stimulation as previously demonstrated.^[5]^ On the other hand, the addition of CDVs derived from wild type C2C12 muscle cells (+CDVs) shortly after plating (8 h) was sufficient to reverse the inhibition of cell growth by magnetic field stimulation in the TRPC1-knockdown clones and, moreover, commenced to reveal a positive trend for magnetically-stimulated myoblast expansion, where the difference between the groups (+/-CDVs) in the TRPC1 knock down clones (c60, c61) reached significance in the pooled data (Figure 5B). Moreover, the capacity of CDVs to rescue magnetic responsiveness tended to be greatest with CDVs derived from wild type myoblasts rather than myotubes (Figure 5B), aligning with previous evidence indicating that TRPC1 is most responsive to magnetic stimulation during the early myoblast stage of myogenesis.^[5]^

**Figure 5.**
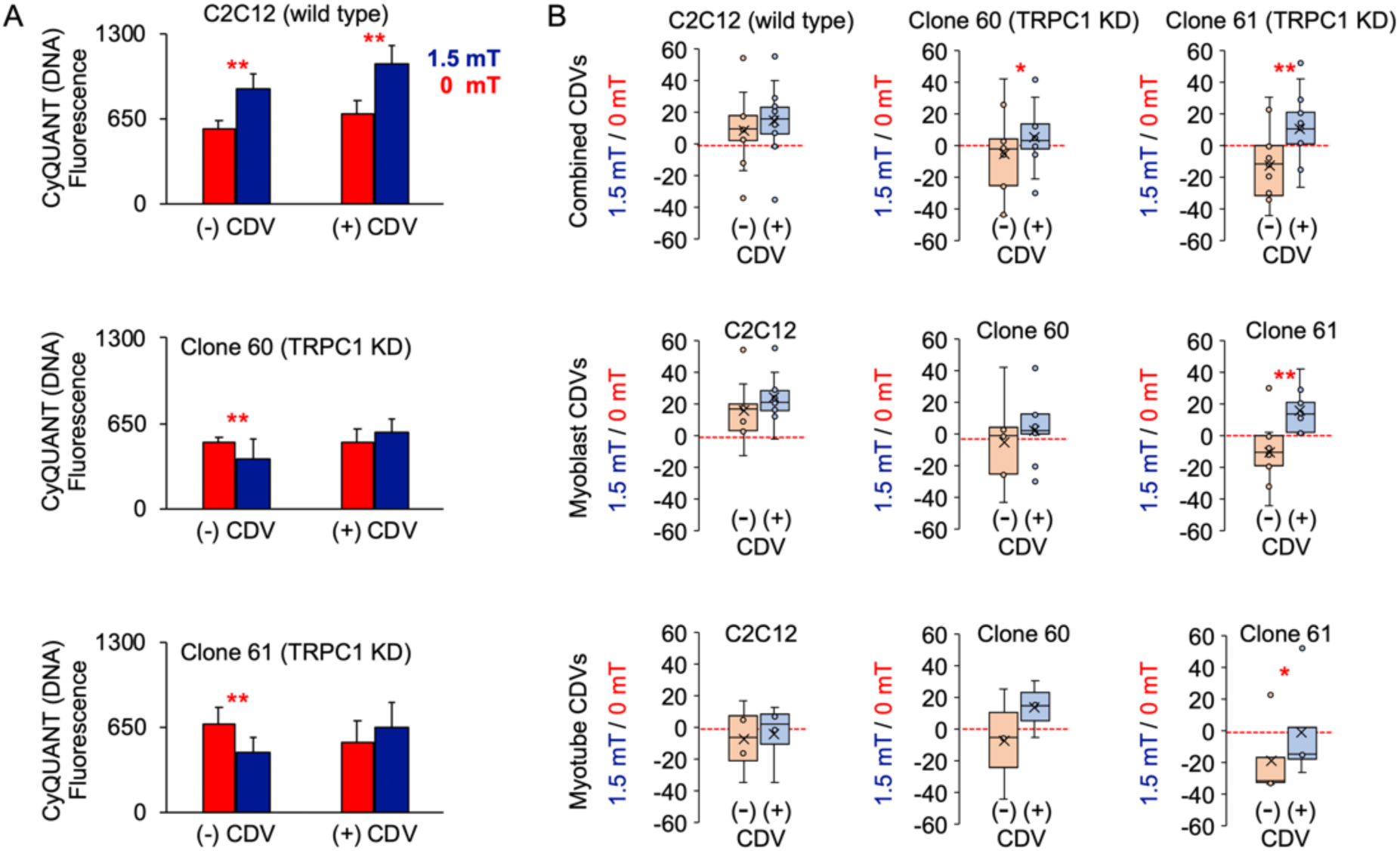
CDV-meditated rescue of magnetoreception in TRPC1 knockdown cells. **A**) Representative magnetically-stimulated proliferative responses of C2C12 wild type (top), c60 TRPC1 knockdown (KD; middle) and c61 TRPC1 knockdown (KD; bottom) myoblast cultures with (+CDV) and without (−CDV) the addition of CDVs derived from C2C12 wild type cultures as indicated. CDVs were added to myoblast cultures 8 h post-cell seeding and 16 h before magnetic exposure at an amplitude of 1.5 mT for 10 min as previously described.^[5]^ Culture DNA content (Cyquant) was measured 24 h after magnetic exposure. Blue and red bars indicate magnetically-exposed and non-magnetically exposed control samples, respectively (\*\**P* < 0.01 with regards to respective 0 mT). n = 12 wells per condition. **B**) Distributions of magnetically-induced proliferation responses (1.5mT/0mT) for wild type C2C12 (left) and CRISPR/Cas9 knockdown C2C12 clones, c60 (middle) and c61 (right) with (blue) and without (orange) the addition of wild type CDVs derived from either myoblasts (middle), myotubes (bottom) or both combined (top). Per condition, the data depicted represents the means of 12 independent experiments (biological replicates), each representing the means of n=10-12 technical replicates. ** and * represent *P* < 0.01 and < 0.05, respectively, pairwise for 0 mT vs 1.5 mT (**A**) or -CDV vs +CDV (**B**).

### 2.7. Functional rescue of myoblast respiratory capacity

Mitochondrial respiration stimulates myoblast proliferation and requires the participation of TRPC1.^[5,22]^ In this respect, low-amplitude PEMFs have been shown to specifically activate TRPC1 and hence, promote myogenesis and the establishment of mitochondrial survival adaptations.^[5]^ Accordingly, myoblasts that had been genetically-engineered to exhibit reduced levels of TRPC1 exhibited a loss of both proliferative and mitochondrial responses to PEMFs. Here we show that mitochondrial magnetic responses could also be partially recovered in such TRPC1-deficient myoblasts upon the addition of wild type CDVs **(Figure 6)** and that the degree of rescue, moreover, is greatest in a mutant cell line (c61) that had previously been shown to possess the greatest depression in basal TRPC1 expression level.^[5]^ The sum of our results thus indicates that TRPC1 channels are necessary and sufficient for magnetoreception that can be conferred to TRPC1-deficient cells with the addition of muscle-derived CDVs.

**Figure 6.**
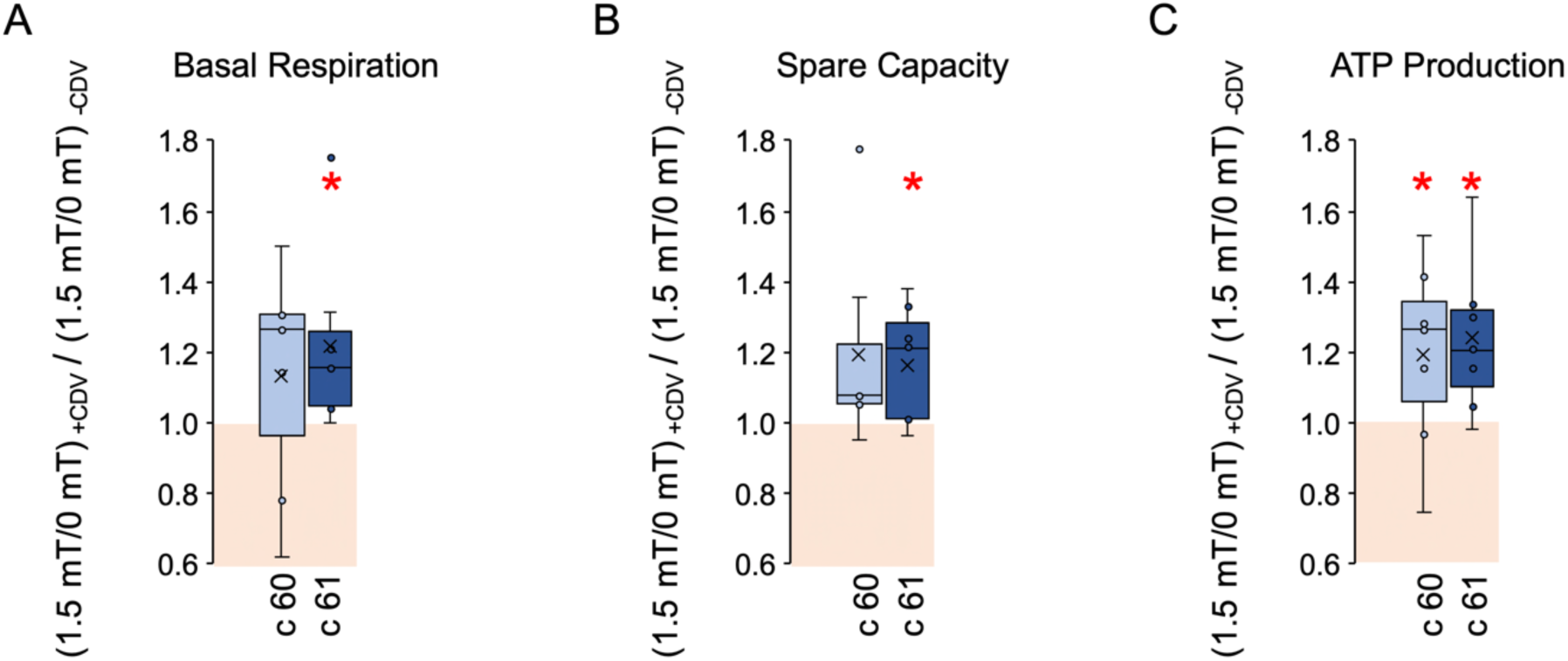
Relative changes [(1.5mT/0mT) plus wt CDVs / (1.5mT/0mT) minus wt CDVs] in PEMF-induced oxygen consumption rate with the use of wild type CDVs. Changes in PEMF-induced Basal Respiration (**A**), Spare Capacity (**B**) and ATP Production (**C**) for CRISPR/Cas9 knockdown C2C12 clones, c60 (light blue) and c61 (dark blue) following the administration of CDVs (+CDVs) relative to untreated cells (−CDVs). Red shaded boxes indicate areas of no or negative change in PEMF response following CDV addition. Data represents the means of 7 independent experiments (biological replicates), each representing the means of technical triplicates. (\**P* < 0.05 relative to untreated cells of the same origin).

## 3. Discussion

Previously published protocols for producing cell-derived vesicles have rendered important information about cellular signal transduction as well as revealed a provocative potential for such platforms to serve as biological-based sensors or as vehicles for drug delivery.^[6–9,54,55]^ Here, we reveal a novel potential of CDVs as biological sensors for temperature (menthol-sensitivity) or magnetic fields and furthermore demonstrated that CDVs possess the capacity to recover sensory transduction of developmental importance for potential therapeutic use in regenerative medicine. Importantly, these signaling and developmental attributes of CDVs can be ascribed to their expression of TRP channels associated with the surface membrane.^[19]^

We provide an in-depth characterization of CDVs derived from C2C12 murine muscle cells, including electron microscopic imaging as well as proteomic and lipidomic analyses. Comparison of our CDV lipid analysis to recently published data obtained from mechanically-derived CDVs ^[16]^ indicates that the procedure for deriving vesicles greatly influences vesicle composition. Here, employing a modified version of the established cytochalasin B protocol, the relative abundancies for PC and PE were substantially lower and those for SM and PS substantially higher than that observed for purely mechanically-derived vesicles, suggesting greater plasma membrane enrichment in our samples. Accordingly, an enrichment in membrane-associated proteins was also rendered by our cytochalasin B protocol. The CDVs generated in this report also exhibited appreciable membrane integrity, demonstrating the capacity to retain soluble intracellular agents and enzymes (e.g. esterases and loaded calcium dyes; *cf.* Figure 4) present within the cell at the time of vesiculation, indicating minimal membrane disruption. Nonetheless, to what extent extra-plasmalemmal compartments may have contributed to the CDV liposome mixture and influence their sensory performance merits further investigation.

TRPC1 promotes myogenesis by enhancing adaptive mitochondrial responses,^[5]^ whereas reductions in TRPC1 expression level are associated with slowed myoblast expansion ^[5,22]^ and attenuated mitochondrial function.^[5]^ Magnetically-induced TRPC1 channel activation hence sets into motion mitochondrial adaptations governing cell survival, in accordance with findings that mitochondrial calcium uptake is critical in supporting oxidative respiration and survival adaptations.^[56]^ By contrast, in quiescent myoblasts characterized by depressed TRPC1, magnetic fields instead inhibit cell growth and attenuate mitochondrial respiration,^[5]^ paralleling the results reported here (Figures 4-6) and suggesting moreover, that TRPC1 exerts a protective effect over mitochondria from being overstimulated by magnetic fields, or otherwise. TRPC1 is hence a key member of the macromolecular complex conferring magnetoreception as well as upstream of mitochondrial functional adaptations to magnetic stimulation. Notably, rescue of magnetic sensitivity by wild type CDVs was most pronounced in the TRPC1 knockdown mutant, c61 (Figures 4-6), which had been previously shown to exhibit the strongest suppression of TRPC1 expression.^[5]^ The nature of the effect exerted by magnetic fields is thus an indication of cellular oxidative/inflammatory status that, in turn, reflects TRPC1 channel expression. Now, with the new understanding that non-invasive magnetic fields can be employed as an adaptive stimulus targeting a TRPC1-mitochondrial axis via a process of magnetic mitohormesis, the possibility exists of reinstating adaptive mitohormesis in senescent or aged tissues with the administration of “proliferative” CDVs in conjunction with magnetic field therapy. Our future experiments will examine the capability of CDVs from proliferative myoblasts to rescue magnetic mitohormesis in quiescent cells, *cf.* ^[5]^.

The function of the cellular secretome is responsive to mitochondrial activation ^[57]^ and has recently been shown to be stimulated by the same magnetic field paradigm employed in this study.^[58]^ Cell senescence moreover, is characterized by a senescence associated secretory phenotype (SASP) ^[2–4]^ that is capable of inducing a contagious state of senescence to neighboring cells and tissues.^[59]^ Therefore, given the enrichment of the healthy muscle secretome for diffusible factors, collectively known as myokines, possessing systemwide therapeutic and regenerative properties,^[60–62]^ the possibility of activating, reinstating or fortifying the myokine response in muscle with CDVs derived from proliferative muscle cells in combination with magnetic field therapy merits further consideration.

Given the recognized requirement of TRPC1 for skeletal muscle development ^[63–65]^ in combination with the importance of skeletal muscle for overall human health and metabolic stabilization,^[60–62]^ methods of reimplementing TRPC1 expression would be of importance during muscular aging or senescence. Additionally, TRPA1 has also been implicated in lifespan extension in *C. elegans* ^[66]^ due to its capacity to reduce age-related oxidative stress.^[67]^ Moreover, menthol has been shown to induce mitochondrial respiration and adaptations in human adipocytes via the activation of menthol-sensitive TRP channels.^[51]^ Therefore, given that both TRPC1 ^[5]^ and TRPA1 ^[67]^ have been linked to mitochondrial function and the accepted correlation between mitochondrial function and longevity,^[68,69]^ the future deployment of delivery systems for these two TRP channels would have clear practical and clinical implications for treating muscle loss or in rescuing states of mitochondrial dysfunction in cases of frailty, advanced age, critical systemic inflammation or post-trauma. Lastly, similar CDV platforms may also be ultimately exploited as magnetically-activated calcium-dependent drug delivery systems given targeted genetic engineering intervention.

At a fundamental level our detailed CDV compositional analyses provide a first-stage database for use by future investigators to target identified molecules and signaling cascades for ultimate exploitation as well as to assist in the development of other CDV paradigms as alternatives to cell-based biosensors. Moreover, we also set the stage for the future development of CDV platforms for the reinstatement of temperature or magnetic mitohormesis in aged or senescent cells or tissues or for optimization of tissue engineering endeavors.

## 4. Experimental Section

### Cell culture and cell-derived vesicles (CDV) production

Otherwise stated, all reagents for cell culture experiments were purchased from Thermo Fischer Scientific, Switzerland. Murine skeletal muscle myoblasts (C2C12; ATCC, USA) were cultured in DMEM containing glucose (4.5 g L^-1^) and sodium pyruvate (1 g L^-1^) and supplemented with 5% fetal bovine serum (FBS) and L-glutamine (2 mmol L^-1^) in a humidified atmosphere at 37 °C and 7% CO_2_. Cell confluence was kept between 20% and 40% to prevent premature differentiation and maintain TRPC1 expression at highest levels.^[5]^ For myotube differentiation, cells were grown to 80% confluence and induced to fuse with the provision of growth medium supplemented with only 2% horse serum for 2 days.

For cell-derived vesicle (CDV) production, myoblasts in 75 cm2 culture flasks were washed once with PBS pH 7.4, incubated for 30 s with trypsin at room temperature (TrypLE express), and then incubated in serum-free RPMI 1640 medium supplemented with cytochalasin B (10 µmol L^-1^; Sigma, Switzerland) for 15 min. The culture flask was mounted onto an incubation shaker (KS 4000 i control, IKA, Germany) set to 300 rpm at 37 °C to increase vesicle formation yield during the incubation with cytochalasin B. The supernatant was subsequently collected and centrifuged at 700 g for 5 min to separate cells and cell debris from the CDV solution. The supernatant was stored at 4 °C until use.

### DLS measurements, liposome extrusion, and osmolarity measurements

Dynamic light scattering (DLS) analysis was used for the size characterization of the CDV samples. Measurements were conducted with a DLS system (Zetasizer 3000HS, Malvern Panalytical Ltd, U.K.) at 25 °C. CDVs were extruded in two steps: first by a polycarbonate membrane with 400 nm diameter pores (11 repetitions), then by a polycarbonate membrane with pores of 200 nm diameter (21 repetitions), according to the manufacturer’s instructions (Avanti Polar Lipids, USA; membranes were purchased from Sigma, Switzerland). Osmolarities of CDV hosting solutions were determined with an osmometer (Osmomat 3000basic; Gonotec, Germany) to assure that changes in osmolarity were ≤ 5% (Table S3).

### Electron microscopy

Sample quality was first assessed by transmission electron microscopy (TEM) of negatively stained samples. Crude samples (non-extruded) were concentrated at 14100 g for 30 min and re-suspended at 32-fold initial concentration prior to imaging (Figure S1A). For cryo-TEM, CDVs were produced and extruded as described above and characterized in their hydrated state in a Tecnai F20 cryo-TEM (FEI, USA). 300-mesh lacey carbon-coated copper grids (Quantifoil, Germany) were glow discharged (Emitech K100X, GB) for 30 s prior to application of 3 µl of sample solution onto the grids in a Vitrobot Mark II (FEI, USA), and the excess of the dispersion was removed by controlled blotting. A mixture of liquid ethane/propane was used for sample vitrification. The grids were then transferred on a Gatan cryo-holder within the microscope and kept at -180 °C during observation. Micrographs were recorded under low dose conditions (< 500 e^-^ nm^-2^) using a 4k x 4k Gatan CCD camera at 200 kV acceleration voltage in bright field mode. To analyze bulk sample organization freeze-fractured samples were imaged by cryo-SEM. CDVs were produced as described above (non-extruded), enriched by an additional 5-fold, and 12 µl of the desired emulsions were sandwiched between two pre-cleaned and hydrophilized 6 mm aluminum planchettes. The samples were frozen in a high-pressure freezer HPM 100 (Bal-Tec/Leica, Vienna) and stored in liquid nitrogen. The frozen samples were subsequently fractured under high vacuum in a pre-cooled freeze-fracture device BAF 060 at -130 °C (Bal-Tec/Leica). Fractured samples were then coated with 2.5 nm of tungsten at a deposition angle of 45° followed by an additional tungsten coating of 2.5 nm at 90° and afterwards transferred to a pre-cooled SEM (−120 °C) (Zeiss Gemini 1530, Oberkochen) for imaging at 2 kV with an in-lens detector.

### Proteome measurements

Total protein content for LC-MS/MS measurements was determined by sodium dodecyl sulfate polyacrylamide gel electrophoresis (SDS-PAGE; PhastSystem, GMI, USA). One milliliter of CDV sample was pelleted at 14100 g for 30 min and re-suspended in 8 µl phosphate buffered saline (PBS; pH 7.4; Thermo Fischer Scientific, Switzerland). After sonication for 10 min, 2 µl of 5x SDS-loading buffer was added to the sample, followed by heating at 85 °C for 5 min. A dilution series was prepared with 1x SDS-loading buffer following with 2 µl of sample being loaded into each lane of a 12.5% SDS gel. Image analysis of the gel was done by ImageJ and quantification extracted from the 1:2 sample dilution (Figure S4). For proteome analysis CDV samples were pooled from 10 T75 flasks, resuspended in PBS pH 7.4, and concentrated 50-fold to yield an approximate total protein content of 300 µg.

For mass spectrometry protein analysis, CDVs were resuspended in lysis buffer containing SDS (2%), Tris (100 mmol L^-1^) and DTT (30 mmol L^-1^). Samples were then precipitated using trichloroacetic acid (TCA) to a final concentration of 20% and incubated on ice for 30 min, followed by centrifugation for 10 min at 14000 rpm. The pellet was washed twice with ice-cold acetone and resuspended in urea (8 mol L^-1^) containing 10% (w/v) RapiGest (Waters). Proteins were then reduced using TCEP (5 mmol L^-1^; tris(2-carboxyethyl)phosphine) for 30 min at RT and subsequently alkylated using iodoacetamide (10 mmol L^-1^) for 30 min at RT in the dark. Samples were diluted to a concentration of urea of 2 mol L^-1^ and digested overnight with the addition of 2 µl sequencing-grade trypsin (Promega). After digestion, the peptide mixture was acidified by adding 0.1% formic acid, incubated at 37 °C for 30 min and centrifuged at 20000 g for 10 min. The supernatant was desalted on C18 UltraMicroSpin columns (The Nest Group) according to the manufacturer’s instructions and dried in a SpeedVac concentrator (Thermo Scientific, Rockford, USA). Peptides were reconstituted in acetonitrile (2%), formic acid (0.1%), and separated by reversed-phase chromatography on a high pressure liquid chromatography (HPLC) column (75 µm inner diameter; New Objective, Woburn, USA), packed in-house with a 15 cm stationary phase (Magic C18AQ, 200 Å, 1.9; Michrom Bioresources, Auburn, USA) and connected to an EASY-nLC 1000 instrument combined with an autosampler (Proxeon, Odense, Denmark). The HPLC was coupled to an Orbitrap Fusion spectrometer equipped with a nanoelectrospray ion source (Thermo Scientific, Rockford, USA). Peptides were loaded onto the column with 100% buffer A (99.9% H_2_O, 0.1% formic acid) and eluted at a constant flow rate of 300 nL min^-1^ for 50 min with a linear gradient from 6-25% buffer B (99.9% acetonitrile, 0.1% formic acid), followed by 15 min from 26-35% buffer B and a washing step with 90% buffer B. Mass spectra were acquired in a data-dependent manner with a 3 s cycle time. In a standard method for medium to low-abundant samples high-resolution MS1 spectra were acquired @ 120000 resolution (automatic gain control target value 2×10^5^) to monitor peptide ions in the mass range of 395–1500 m z^-1^, followed by HCD MS/MS scans @ 30000 resolution (automatic gain control target value 1×10^2^). To avoid multiple scans of dominant ions, the precursor ion masses of scanned ions were dynamically excluded from MS/MS analysis for 60 s. Single charged ions and ions with unassigned charge states or charge states above 6 were excluded from MS/MS fragmentation. For data analysis, SEQUEST software v27.029 was used to search fragment ion spectra for a match to fully tryptic peptides without missed cleavage sites from a protein database, which was composed of the Mus musculus proteome (SwissProt), various common contaminants and sequence-reversed decoy proteins. The precursor ion mass tolerance was set to 20 ppm. Carbamidomethylation was set as a fixed modification on all cysteines. The PeptideProphet and the ProteinProphet tools of the Trans-Proteomic Pipeline (TPP v4.6.2) were used for the probability scoring of peptides spectrum matches 30, and inferred protein identifications were filtered to reach an estimated false-discovery rate of ≤1%. The same sample was reinjected and selected proteotypic peptides mapping to TRPC1, ORAI1, TRPA1, TRPV1, and TRPV2 were monitored using Parallel-Reaction Monitoring as previously described.^[70]^ Channel structures including highlighted peptides targeted for mapping TRPC1, ORAI1, and TRPA1 are plotted in Figure S5.

### Gene Ontology Analysis

Protein associations to cell components, molecular function, and biological processes were retrieved from the panther gene ontology tool (www.pantherdb.org). CDV data was compared to C2C12 data published by Deshmukh et al.,^[31]^ which provides the most substantial and detailed proteome analysis of C2C12 cultures to date.

### Lipid analyses

CDV samples for lipid analysis were prepared identically to samples for protein analysis. Cell samples were collected from a T25 cm2 culture flask, washed once with PBS (pH 7.4), and the cell pellet was frozen in liquid nitrogen. Samples were stored at −80 °C until subsequent sample preparation for analysis.

Lipid extraction was performed as described previously ^[71]^ with some modifications. One milliliter of a mixture of methanol:MTBE:chloroform (MMC) 1.33:1:1 (v/v/v) was added to 20 µl of sample. The MMC was fortified with the SPLASH mix of internal standards (Avanti Lipids) and 100 pmol mL^-1^ of the internal standards: d7-sphinganine (d18:0), d7-sphingosine (d18:1), dihydroceramide (d18:0:12:0), ceramide (d18:1/12:0), glucosylceramide (d18:1/8:0), sphingomyelin (18:1/12:0), and 50 pmol mL^-1^ d7-sphingosine-1-phosphate. After brief vortexing, the samples were continuously mixed in a Thermomixer (Eppendorf) at 25 °C (950 rpm, 30 min). Protein precipitation was obtained after centrifugation for 10 min at 16000 g and 25 °C. The single-phase supernatant was collected, dried under N_2_, and stored at –20 °C until analysis. Before analysis, the dried lipids were redissolved in 100 µl methanol:isopropanol (1:1; v/v).

Liquid chromatography was conducted as described previously ^[72]^ with minor modifications. The lipids were separated using C30 reverse phase chromatography. A transcend TLX eluting pump (Thermo Scientific) was used with the following mobile phases: (A) acetonitrile:water (6:4) with ammonium acetate (10 mmol L^-1^) and 0.1% formic acid and (B) isopropanol:acetonitrile (9:1) with ammonium acetate (10 mmol L^-1^) and 0.1% formic acid. A C30 Accucore LC column (Thermo Scientific) with the dimensions 150 mm x 2.1 mm x 2.6 µm (length x internal diameter x particle diameter) was used. The following gradient was used with a total flow rate of 0.26 ml min^-1^, whereby the mobile phase B was changes as the following: (i) 0.0-0.5 min (isocratic 30% B); (ii) 0.5-2 min (ramp 30-43% B); (iii) 2.10-12.0 min (ramp 43-55% B); (iv) 12.0-18.0 min (ramp 65-85%); (v) 18.0-20.0 min (ramp 85%-100% B); (vi) 20-35 min (isocratic 100% B); (vii) 35-35.5 min (ramp 100-30% B) and; (viii) 35.5-40 min (isocratic 30% B).

The liquid chromatography was coupled to a hybrid quadrupole-orbitrap mass spectrometer (Q-Exactive, Thermo Scientific). Data-dependent acquisition was conducted with positive and negative polarity switching. A full scan was acquired from 220-3,000 m z^-1^ at a resolution of 70000 and an automatic gain control (AGC) target of 3 x 10^6^, while data-dependent scans (top 10) were acquired using normalized collision energies (NCE) of 25 and 30 at a resolution of 17500 and an AGC target of 1 x 10^5^.

Identification of the lipids was achieved using 4 criteria: (1) high accuracy and resolution with a m z^-1^ accuracy within 5 ppm shift from the predicted mass and a resolving power of 70000 at 200 m z^-1^; (2) isotopic pattern fitting to expected isotopic distribution; (3) comparing the expected retention time to an in-house database and; (4) the fragmentation pattern matching to an in-house experimentally-validated lipid fragmentation database.

Quantification was done using single point calibration by comparing the area under the peak of each ceramide species to the area under the peak of the internal standard. Quality controls using a mixture of all samples were used in 4 concentrations (1X, 0.5X, 0.25X, and 0.125X). Triplicates on the QCs were measured and the CV% for each of the lipids reported was below 20%. The following lipid classes were identified in the current study: aclycarnitites, phospholipids (phosphatidylcholines, phospatidylethanolamines, phosphatidylinositols, phosphatidylglycerols, and phosphatidyserines), sphingolipids (sphingoid base phosphates, ceramides, deoxyceramides, monohexosylceramides, and sphinogmeylins), and glycerolipids (diaclyglyerols, triacylglyercerols). Mass spectrometric data analysis was performed in Treacefinder software 4.1 (Thermo Scientific) for peak picking, annotation, and matching to the inhouse fragmentation database.

### Microfluidic chip, CDV immobilization, and microscopy

The microfluidic chip design, fabrication, and vesicle immobilization procedure have been previously described.^[28,50]^ Briefly, poly(dimethyl)siloxane (Sylgard 184, Dow Corning, USA) was mixed at a ratio of 10:1 (pre-polymer : curing agent), poured onto the master mold, and cured over night at 80 °C. The finalized polymer chips were then bonded onto a glass cover slip (#1.5, Menzel Gläser, Germany) after micro-contact printing of biotinylated bovine serum albumin (bBSA) patterns.

All non-printed surfaces were immediately blocked with BSA followed by formation of the molecular cholesterol linkers for vesicle capture (bBSA:avidin:bBSA-PEG-cholesterol). CDVs were immobilized on these surface patches directly before measurement in serum-free RPMI 1640 at 37 °C controlled by a custom-made stage incubation system.^[73]^ CDV membrane integrity and cytosolic content were validated with calcein-AM (1 µmol L^-1^ final concentration; 1 mmol L^-1^ stock solution in DMSO; Thermo Fischer Scientific, Switzerland) membrane partitioning, enzymatic cleavage by endogenous esterases, and subsequent retention within CDVs. Calcein-AM was administered in serum-free RPMI 1640 medium for 10 min followed by 10 min of solution exchange within the microfluidic channel. Imaging of calcein fluorescence occurred by epifluorescence microscopy using a 60X water immersion objective (NA 1.2), a halide arc lamp excitation source (X-Cite,120 PCQ, Lumen Dynamics, Canada) and appropriate bandpass filter sets (ex. 470/40, 500 dichroic, em. 525/50) on an inverted microscope (IX70, Olympus, Germany). Printed protein patterns were imaged by monitoring Texas red fluorescence (ex. 546/12, 560 dichroic, em. 607/80).

To quantify the effect of menthol on calcium influx into the CDVs, the vesicles were loaded with Fluo-4 AM (1 µmol L^-1^ final concentration; 1 mmol L^-1^ stock solution in DMSO; Thermo Fischer Scientific, Switzerland) for 10 min followed by incubation for another 10 min. Hosting fluid was exchanged and menthol was administered at 500 µmol L^-1^ final concentration (500 mmol L^-1^ stock solution in ethanol (analytical grade); Sigma, Switzerland) in serum-free RPMI 1640. Imaging commenced immediately upon initiation of fluid flow to monitor menthol-induced calcium influx. The data was generated from 3 separate microfluidic channels, 2 channels with menthol and 1 channel without menthol to serve as control, using CDVs derived from the same batch of cells. The only difference in experimental conditions was the presence/absence of menthol. The Fluo 4 signal was monitored with the same setup used for the calcein-AM test. The Fluo-4 signal intensity represented the mean fluorescence intensity of a CDV-immobilized region being imaged minus background-subtraction from areas of the microfluidic channel not containing immobilized CDVs. Either areas could be easily differentiated from the other due to presence of Texas red-conjugated avidins deposited as part of the immobilization functionalization protocol.

### CDV-cell fusion experiments

Myoblast cultures after 48 h of growth were stained with the lipophilic membrane dye R18 (ThermoFischer Scientific, Switzerland) at a final concentration of 1 µmol L^-1^ in PBS for 10 min immediately prior to CDV preparation as described before (see *Cell culture and cell-derived vesicles (CDV) production*). The resulting CDV pellet was resuspended in serum-free RPMI 1640 at half of the original volume. Recipient myoblasts for ultimate CDV fusion experiments were plated into multi-well slides (Ibidi, Germany) the day before CDV preparation. On the day of the experiment, the culture medium of the recipient myoblasts was replaced with 200 µl of the stained CDV-suspension and incubated for 30 min at 37 °C and 7% CO_2_. The solution was then exchanged for 200 µl serum-free DMEM medium without phenol red in preparation for imaging with a confocal microscope (Visitron, Germany) equipped with a 60X/1.2 NA water immersion objective, a spinning disk confocal unit (CSU-W1), and a live cell imaging chamber.

### CRISPR/Cas9 TRPC1 exon 1 deletion in C2C12 muscle cells

Myoblast clones 60 (TRPC1 knockdown by ∼40%) and 61 (TRPC1 knockdown by ∼60%) were previously described ^[5]^. Briefly, candidate sgRNAs (single-guide RNA) targeting the ATG start site in exon 1 of the mouse TRPC1 (NC_000075.6) were generated using CRISPR Design Tool (http://crispr.mit.edu). Two candidate sgRNAs of 20 nucleotides in length were cloned into PX458 (pSpCas9-P2A-GFP) and PX459 (pSpCas9-P2A-Puro), respectively. The fortuitous presence of several lower efficiency alternative 5’ upstream non-AUG translational start sites allowed functional depression (knockdown) of TRPC1 in clones 60 and 61 without the deleterious developmental side effects of complete silencing of TRPC1.

### Pulsed Electromagnetic Field (PEMF) Device

The PEMF device used in this study has been previously described.^[5,20,58]^ All samples were exposed once for 10 min at a field amplitude of 1.5 mT. All PEMF-treated samples were compared to time-matched control samples (0 mT) that were manipulated in exactly the same as experimental samples, including placement into the PEMF-generating apparatus for the designated time, except that the apparatus was not set to generate a magnetic field.

### Calcium Imaging

Myoblasts or myotubes from C2C12 wild type or C2C12-derived TRPC1 CRISPR/Cas9 knockdown clones (C60 and C61) were incubated with Calcium Green-1, AM (50 µg in 100 µl DMSO; Invitrogen, Singapore) for 30 min at 37 °C in the dark. Cell cultures were then rinsed with PBS and CDVs prepared as described previously (see *Cell culture and cell-derived vesicles (CDV) production*). CDVs were resuspended in 200 µl of FluoroBrite medium (Gibco, Singapore) and exposed to Pulsed electromagnetic fields (PEMFs) at 0 mT or 1.5 mT for 10 min. Post-magnetic exposure, the CDVs were seeded into black bottomed 96-well polystyrene plates (Thermo Fisher, Singapore, Cat no. 137101) and the calcium levels measured using a Cytation5 (BioTek, Singapore) plate reader at 500 nm excitation and 535 nm emission wavelength with bandwidths set to 9 nm excitation and 20 nm emission wavelengths.

### Cellular proliferation assay (DNA content)

C2C12, C20, C60 and C61 cells were seeded into 96-well plates (Thermo Fisher, Singapore, Cat. no. 167008) at a density of 1000 cells/well in replicates of 10 wells. Cultures of myoblasts were administered 50 µl of CDVs generated from either C2C12 myoblast or myotube cultures, 8 h post cell seeding. The CDV-impregnated dishes were exposed to PEMF 24 h post-seeding at 0 mT or 1.5 mT for 10 min. 24 h following PEMF exposure the cells were incubated with 100 µl of CYQUANT direct reagent (Thermo Scientific, Singapore) for 2 h before analysis at 480/535 nm using a Cytation5 plate reader (BioTek, Singapore).

### Metabolic Flux Analysis (oxygen consumption rate)

Myoblast oxygen consumption rate was measured as previously described.^[5]^ Briefly, C2C12 myoblasts were seeded into Agilent Seahorse XFe24 plates in growth media (GM) at a density of 20,000 cells/well. 150 µl of CDVs were added to the adherent cells 4 h post seeding and incubated overnight before PEMF exposure. Following PEMF exposure for 10 min myoblasts were allowed to stabilize for 2 h in a standard humidified incubator at 37°C and 5% CO_2_. In preparation for mitochondrial assessment, the culture media was replaced with XF Base medium (Agilent Technologies, Inc., USA) consisting of glucose (10 mmol L^-1^), sodium pyruvate (1 mmol L^-1^), and glutamine (2 mmol L^-1^), pH 7.4 and equilibrated in a non-CO_2_ environment at 37°C for 60 min as per manufacturer’s protocol. Cellular respiratory capacity was assayed using the XF Cell Mito Stress Test Kit (Agilent Technologies, Inc., USA). The assay consists of repeated cycles of 3-min mix, 2-min wait and 3-min measurement cycles for baseline and after each injection of Oligomycin (1 µmol L^-1^), FCCP (1 µmol L^-1^) and Rotenone + Antimycin A (0.5 µmol L^-1^). Data analysis was done using Seahorse Wave Desktop Software (Agilent Technologies, Inc., USA).

## Supporting Information

Supporting Information is available in separate files or from the authors.

## Supporting information

Kurth_et_al_2020_bioRxiv_SI

Proteotype_CDV_C2C12

## Acknowledgements

The authors would like to thank the Hilvert laboratory at ETH Zurich for support with SDS page, S. Krämer for use of the DLS system, and S. Handschin from the ScopeM facility for electron microscopy. The authors also acknowledge Zac Goh (iHealthtech, National University of Singapore) for the graphical depiction of our basic findings shown in Figure 1 and 4A. P.S.D. acknowledges financial support from the European Research Council (ERC consolidator grant No. 681587). B.W. and MvO would like to acknowledge the ETH (grant ETH-25 15-2) and the Swiss National Science Foundation (grant 31003A_160259) for funding support. A.F.O. would like to acknowledge the Institute for Health Innovation & Technology, iHealthtech, at the National University of Singapore for helping support this study. A.F.O. would also like to thank the Lee Foundation, Singapore (R-176-000-243-731) and National University of Singapore for financially supporting this research.

