## Supplementary material for "Cell-Derived Vesicles as TRPC1 Channel Delivery Systems for the Recovery of Cellular Respiratory and Proliferative Capacities": Kurth_et_al_2020_bioRxiv_SI

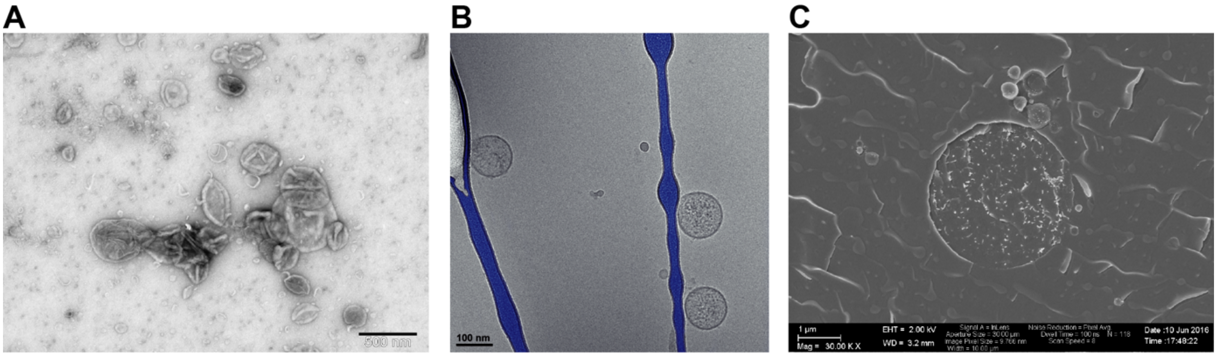

**Figure S1.** (A) Heavy metal labeled cryo-TEM image of cell-derived vesicles (CDVs) generated from C2C12 muscle cells. (B) Cryo-TEM image from an analogous CDV sample. The granular intraliposomal content is clearly distinguishable from the more homogeneous hosting solution background. Areas of the carbon-free grid are colored in blue for illustrative purposes. (C) Freeze fracture SEM image of a large CDV cleaved through its lumen upon sample fracture and revealing a predominance of granular content.

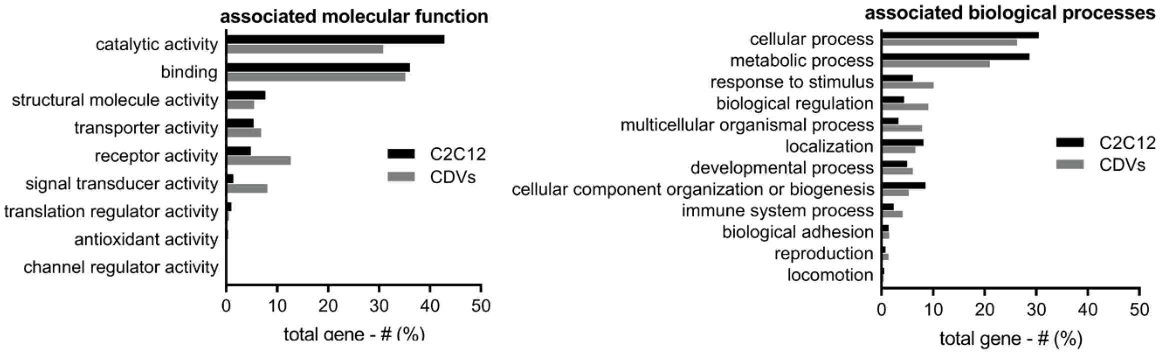

**Figure S2.** Molecular functions (left) and biological processes (right) associated with the identified proteins from C2C12 cultures and CDV samples. Molecular functions that are characteristic of the plasma membrane were represented in relatively higher abundances in CDVs samples.

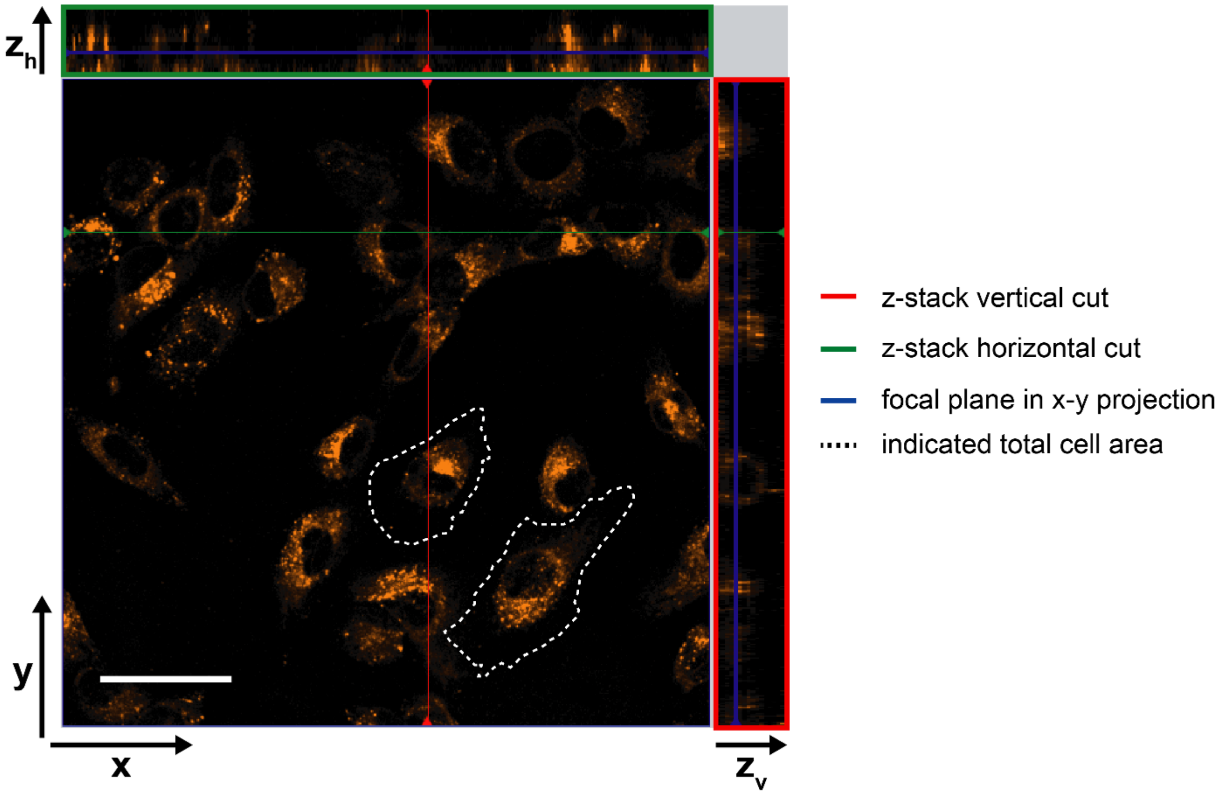

**Figure S3.** Confocal micrograph depicting horizontal and vertical z-projections. The z-stack of the horizontal cut (green) is located above and the vertical cut (red) on the right of the micrograph. The green and red lines in the micrograph delineate the paths of the horizontal and vertical cuts, respectively. The blue lines in the z-projection delineate the focal plane in the z-stack, which is displayed as the x-y confocal projection. The white dotted lines demarcate the discernable peripheries of 2 individual cells. Scale bar: 20 µm.

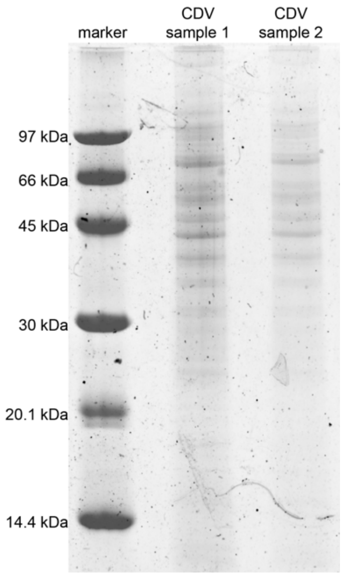

**Figure S4.** SDS-PAGE gel of CDV samples run for the determination of total protein concentration. Samples were diluted 1:1 and 1:2 in 1x SDS loading buffer for samples 1 and 2, respectively. The gel scan was adjusted for brightness and contrast.

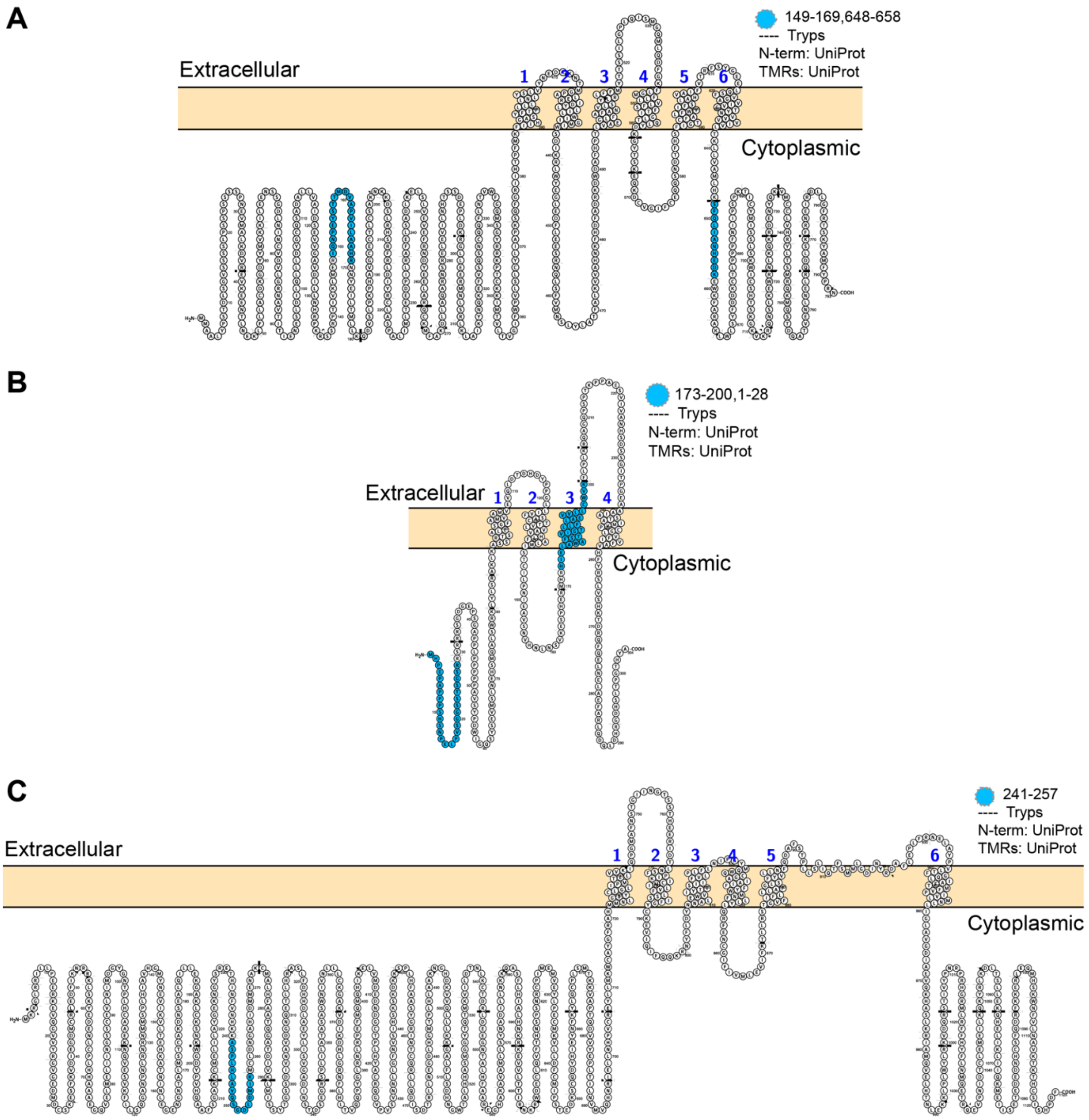

**Figure S5.** Visualization of the channels TRPC1 (A), ORAI1 (B), and TRPA1 (C). Peptides targeted during Parallel Reaction Monitoring are marked in blue. Sequence, topology, and annotations available on Protter (http://wlab.ethz.ch/protter/) via the following links: https://bit.ly/3eTESae (TRPC1), https://bit.ly/2yPdtWl (ORAI1), https://bit.ly/2xatPsf (TRPA1).

**Table S1.** GO cell component enrichment for CDVs over C2C12 cultures derived by the panther GO platform.

| Description | P-value | FDR q-value | Enrichment |
|---|---|---|---|
| filopodium | 6.07E-07 | 9.56E-06 | 1.91 |
| podosome | 3.05E-05 | 3.53E-04 | 1.91 |
| immunological synapse | 1.12E-04 | 1.18E-03 | 1.91 |
| proton-transporting two-sector ATPase complex | 1.12E-04 | 1.17E-03 | 1.91 |
| eukaryotic translation initiation factor 3 complex | 7.91E-04 | 6.58E-03 | 1.91 |
| proton-transporting two-sector ATPase complex, catalytic domain | 7.91E-04 | 6.54E-03 | 1.91 |
| cytosolic large ribosomal subunit | 6.15E-10 | 1.53E-08 | 1.86 |
| ruffle | 4.98E-08 | 9.55E-07 | 1.85 |
| myelin sheath | 4.68E-32 | 4.42E-30 | 1.84 |
| blood microparticle | 1.11E-06 | 1.63E-05 | 1.84 |
| cytosolic small ribosomal subunit | 2.06E-06 | 2.99E-05 | 1.84 |
| ruffle membrane | 2.37E-05 | 2.85E-04 | 1.82 |
| endoplasmic reticulum lumen | 4.35E-05 | 4.88E-04 | 1.82 |
| cortical cytoskeleton | 2.19E-07 | 3.81E-06 | 1.8 |
| axon terminus | 2.64E-04 | 2.51E-03 | 1.8 |
| dendritic shaft | 2.64E-04 | 2.49E-03 | 1.8 |
| neuromuscular junction | 2.64E-04 | 2.48E-03 | 1.8 |
| extrinsic component of plasma membrane | 3.97E-07 | 6.41E-06 | 1.79 |
| cortical actin cytoskeleton | 4.78E-04 | 4.30E-03 | 1.79 |
| presynapse | 4.19E-06 | 5.71E-05 | 1.78 |
| actomyosin | 2.19E-08 | 4.46E-07 | 1.77 |
| stress fiber | 2.23E-07 | 3.78E-06 | 1.76 |
| contractile actin filament bundle | 2.23E-07 | 3.73E-06 | 1.76 |
| terminal bouton | 1.34E-05 | 1.67E-04 | 1.76 |
| extrinsic component of cytoplasmic side of plasma membrane | 1.32E-04 | 1.34E-03 | 1.74 |
| neuron projection terminus | 1.32E-04 | 1.33E-03 | 1.74 |
| actin filament bundle | 1.14E-07 | 2.16E-06 | 1.73 |
| leading edge membrane | 3.82E-06 | 5.27E-05 | 1.73 |
| polysome | 7.02E-04 | 5.92E-03 | 1.71 |
| cell leading edge | 7.02E-04 | 5.88E-03 | 1.71 |
| neuron spine | 3.14E-06 | 4.37E-05 | 1.7 |
| dendritic spine | 5.39E-06 | 7.13E-05 | 1.69 |
| postsynapse | 5.39E-06 | 7.06E-05 | 1.69 |
| focal adhesion | 8.44E-31 | 7.44E-29 | 1.68 |

| | | | |
|---|---|---|---|
| cell-substrate junction | 1.11E-30 | 9.21E-29 | 1.68 |
| cell-substrate adherens junction | 1.95E-30 | 1.52E-28 | 1.68 |
| large ribosomal subunit | 1.18E-08 | 2.48E-07 | 1.68 |
| cluster of actin-based cell projections | 1.57E-05 | 1.93E-04 | 1.68 |
| actin-based cell projection | 2.19E-07 | 3.77E-06 | 1.67 |
| brush border | 2.68E-05 | 3.16E-04 | 1.67 |
| cell periphery | 3.03E-04 | 2.82E-03 | 1.67 |
| endoplasmic reticulum-Golgi intermediate compartment | 5.12E-04 | 4.52E-03 | 1.66 |
| intercalated disc | 8.61E-04 | 7.04E-03 | 1.65 |
| ribosomal subunit | 1.58E-12 | 5.22E-11 | 1.64 |
| postsynaptic specialization | 2.82E-09 | 6.66E-08 | 1.64 |
| postsynaptic density | 2.82E-09 | 6.54E-08 | 1.64 |
| sperm part | 5.35E-05 | 5.95E-04 | 1.63 |
| adherens junction | 1.87E-41 | 3.09E-39 | 1.62 |
| anchoring junction | 3.60E-41 | 5.29E-39 | 1.62 |
| cell-cell adherens junction | 9.30E-21 | 5.59E-19 | 1.62 |
| axon part | 2.13E-08 | 4.40E-07 | 1.62 |
| cell cortex part | 5.35E-07 | 8.53E-06 | 1.62 |
| perikaryon | 5.84E-04 | 5.05E-03 | 1.62 |
| actin filament | 1.45E-04 | 1.46E-03 | 1.61 |
| sarcoplasmic reticulum | 9.57E-04 | 7.72E-03 | 1.61 |
| small ribosomal subunit | 1.51E-05 | 1.87E-04 | 1.6 |
| synaptic vesicle | 2.37E-04 | 2.33E-03 | 1.6 |
| cytosolic part | 1.91E-11 | 5.49E-10 | 1.59 |
| myosin complex | 3.86E-04 | 3.55E-03 | 1.59 |
| synapse part | 5.47E-16 | 2.34E-14 | 1.58 |
| cell cortex | 6.36E-05 | 6.89E-04 | 1.58 |
| lamellipodium | 2.72E-06 | 3.87E-05 | 1.57 |
| exocytic vesicle | 2.51E-04 | 2.44E-03 | 1.57 |
| growth cone | 1.68E-05 | 2.04E-04 | 1.56 |
| intermediate filament | 4.01E-04 | 3.66E-03 | 1.56 |
| ribosome | 1.45E-10 | 3.83E-09 | 1.55 |
| mitochondrial protein complex | 1.92E-07 | 3.39E-06 | 1.55 |
| cell projection membrane | 7.41E-07 | 1.13E-05 | 1.55 |
| extracellular organelle | 2.74E-107 | 3.63E-104 | 1.54 |
| extracellular vesicle | 2.74E-107 | 1.81E-104 | 1.54 |
| extracellular exosome | 3.41E-107 | 1.50E-104 | 1.54 |
| inner mitochondrial membrane protein complex | 2.85E-06 | 4.01E-05 | 1.54 |

| | | | |
|---|---|---|---|
| basolateral plasma membrane | 2.68E-05 | 3.14E-04 | 1.54 |
| proteasome complex | 1.63E-04 | 1.62E-03 | 1.54 |
| endopeptidase complex | 1.63E-04 | 1.61E-03 | 1.54 |
| extracellular matrix | 2.41E-11 | 6.64E-10 | 1.53 |
| membrane region | 5.95E-11 | 1.61E-09 | 1.53 |
| site of polarized growth | 4.19E-05 | 4.73E-04 | 1.53 |
| coated vesicle | 2.54E-04 | 2.45E-03 | 1.53 |
| cytoplasmic region | 2.65E-05 | 3.16E-04 | 1.52 |
| cell-cell junction | 1.10E-20 | 6.30E-19 | 1.51 |
| membrane microdomain | 3.28E-09 | 7.49E-08 | 1.51 |
| membrane raft | 3.28E-09 | 7.36E-08 | 1.51 |
| secretory vesicle | 1.87E-07 | 3.49E-06 | 1.51 |
| actin cytoskeleton | 7.02E-07 | 1.09E-05 | 1.51 |
| endocytic vesicle | 6.10E-04 | 5.17E-03 | 1.51 |
| cell junction | 1.62E-35 | 1.78E-33 | 1.5 |
| perinuclear region of cytoplasm | 7.81E-14 | 2.95E-12 | 1.5 |
| cell body | 1.27E-12 | 4.32E-11 | 1.5 |
| extrinsic component of membrane | 1.02E-05 | 1.28E-04 | 1.5 |
| vesicle | 1.18E-100 | 3.90E-98 | 1.49 |
| apical plasma membrane | 6.05E-06 | 7.77E-05 | 1.49 |
| secretory granule | 6.03E-05 | 6.59E-04 | 1.49 |
| extracellular region part | 7.80E-95 | 2.07E-92 | 1.48 |
| neuron part | 4.92E-26 | 3.25E-24 | 1.48 |
| neuron projection | 3.78E-18 | 2.09E-16 | 1.48 |
| transport vesicle | 8.60E-05 | 9.18E-04 | 1.48 |
| transmembrane transporter complex | 5.43E-04 | 4.76E-03 | 1.48 |
| plasma membrane raft | 5.43E-04 | 4.73E-03 | 1.48 |
| respiratory chain complex | 8.79E-04 | 7.14E-03 | 1.48 |
| cell projection part | 8.10E-17 | 3.83E-15 | 1.47 |
| organelle outer membrane | 1.29E-04 | 1.33E-03 | 1.47 |
| outer membrane | 1.29E-04 | 1.32E-03 | 1.47 |
| Z disc | 3.36E-04 | 3.11E-03 | 1.47 |
| neuronal cell body | 1.18E-08 | 2.51E-07 | 1.46 |
| axon | 7.41E-07 | 1.14E-05 | 1.46 |
| peptidase complex | 5.04E-04 | 4.47E-03 | 1.46 |
| dendrite | 2.41E-07 | 3.99E-06 | 1.45 |
| mitochondrial membrane part | 2.63E-06 | 3.78E-05 | 1.45 |
| synapse | 9.26E-07 | 1.38E-05 | 1.44 |

| | | | |
|---|---|---|---|
| mitochondrial outer membrane | 4.23E-04 | 3.83E-03 | 1.44 |
| plasma membrane region | 4.89E-12 | 1.58E-10 | 1.43 |
| contractile fiber part | 3.50E-05 | 4.03E-04 | 1.43 |
| cell projection | 2.58E-24 | 1.63E-22 | 1.42 |
| receptor complex | 5.03E-04 | 4.49E-03 | 1.42 |
| oxidoreductase complex | 8.13E-04 | 6.68E-03 | 1.42 |
| plasma membrane protein complex | 1.07E-04 | 1.14E-03 | 1.41 |
| organelle lumen | 4.47E-06 | 6.03E-05 | 1.39 |
| membrane-enclosed lumen | 4.47E-06 | 5.97E-05 | 1.39 |
| intracellular organelle lumen | 6.35E-06 | 8.07E-05 | 1.39 |
| membrane protein complex | 2.04E-11 | 5.75E-10 | 1.38 |
| mitochondrial membrane | 2.49E-10 | 6.46E-09 | 1.37 |
| organelle inner membrane | 5.07E-09 | 1.12E-07 | 1.37 |
| intracellular | 5.44E-09 | 1.18E-07 | 1.37 |
| plasma membrane part | 9.07E-18 | 4.62E-16 | 1.36 |
| mitochondrial inner membrane | 3.84E-08 | 7.46E-07 | 1.36 |
| lytic vacuole membrane | 5.98E-04 | 5.13E-03 | 1.36 |
| lysosomal membrane | 5.98E-04 | 5.10E-03 | 1.36 |
| intracellular vesicle | 2.74E-09 | 6.71E-08 | 1.35 |
| cytoplasmic vesicle | 2.74E-09 | 6.59E-08 | 1.35 |
| supramolecular polymer | 1.87E-07 | 3.44E-06 | 1.35 |
| supramolecular complex | 1.87E-07 | 3.40E-06 | 1.35 |
| supramolecular fiber | 1.87E-07 | 3.35E-06 | 1.35 |
| bounding membrane of organelle | 4.43E-10 | 1.13E-08 | 1.34 |
| whole membrane | 2.46E-08 | 4.92E-07 | 1.34 |
| polymeric cytoskeletal fiber | 2.59E-07 | 4.22E-06 | 1.34 |
| cytosol | 7.88E-14 | 2.90E-12 | 1.33 |
| vacuolar membrane | 3.60E-05 | 4.11E-04 | 1.33 |
| mitochondrial part | 1.37E-11 | 4.02E-10 | 1.32 |
| organelle membrane | 6.60E-16 | 2.73E-14 | 1.31 |
| cytoskeleton | 9.84E-13 | 3.42E-11 | 1.31 |
| cell surface | 6.67E-05 | 7.18E-04 | 1.31 |
| cytoskeletal part | 5.72E-12 | 1.80E-10 | 1.29 |
| vacuolar part | 2.61E-04 | 2.51E-03 | 1.27 |
| plasma membrane | 2.81E-17 | 1.38E-15 | 1.26 |
| mitochondrion | 4.09E-16 | 1.81E-14 | 1.26 |
| endoplasmic reticulum part | 5.44E-05 | 6.00E-04 | 1.26 |
| cytoplasmic part | 8.94E-58 | 1.97E-55 | 1.22 |

| | | | |
|---|---|---|---|
| membrane | 3.14E-45 | 5.93E-43 | 1.22 |
| endoplasmic reticulum | 5.40E-06 | 7.00E-05 | 1.2 |
| protein complex | 3.01E-13 | 1.07E-11 | 1.19 |
| cytoplasm | 5.72E-28 | 3.98E-26 | 1.18 |
| non-membrane-bounded organelle | 8.78E-12 | 2.70E-10 | 1.18 |
| intracellular non-membrane-bounded organelle | 8.78E-12 | 2.64E-10 | 1.18 |
| macromolecular complex | 3.70E-15 | 1.49E-13 | 1.16 |
| organelle part | 3.01E-16 | 1.37E-14 | 1.12 |
| intracellular organelle part | 4.18E-15 | 1.63E-13 | 1.12 |
| membrane part | 2.59E-08 | 5.12E-07 | 1.12 |
| organelle | 1.19E-37 | 1.43E-35 | 1.1 |
| membrane-bounded organelle | 6.52E-30 | 4.79E-28 | 1.1 |
| intracellular part | 1.55E-34 | 1.58E-32 | 1.08 |
| intracellular organelle | 5.70E-18 | 3.02E-16 | 1.08 |
| cell part | 6.72E-40 | 8.89E-38 | 1.07 |
| intracellular membrane-bounded organelle | 7.57E-07 | 1.14E-05 | 1.06 |

**Table S2.** Detailed lipid measurement results (LC-MS) of CDVs and C2C12 cultures. Mean values originate from 3 independently measured samples, STD is the standard deviation. Lipid abbreviations are: TG: triglyceride, SM: sphingomyelin, PS: phosphatidylserine, PI: phosphatidylinositol, PE: phosphatidylethanolamine, PC: phosphatidylcholine, LPC: lysophosphatidylcholine, CER: ceramide

| Lipid | CDVs | | C2C12 | |
|---|---|---|---|---|
| *(values in pmol/25µl)* | MEAN | STD | MEAN | STD |
| Cer (d42:3) (d18:1/24:2) | 4.01 | 1.08 | 2.34 | 0.10 |
| Cer (d42:3) (d18:2/24:1) | 4.01 | 1.08 | 2.34 | 0.10 |
| CerG (d40:1) (d18:1/22:0) | 11.77 | 2.85 | 7.60 | 0.50 |
| CerG (d42:1) (d18:1/24:0) | 26.42 | 7.21 | 15.97 | 0.78 |
| CerG (d42:2) (d18:1/24:1) | 30.18 | 5.16 | 21.97 | 1.66 |
| LPC (14:0) | 1.41 | 0.88 | 4.32 | 0.27 |
| LPC (18:1) | 17.23 | 2.61 | 23.60 | 3.01 |
| LPC (24:0) | 15.42 | 1.19 | 13.58 | 0.43 |
| LPC (18:0) | 34.34 | 4.81 | 19.78 | 1.87 |
| LPC (16:0) | 43.45 | 7.64 | 33.88 | 3.30 |

| | | | | |
|---|---|---|---|---|
| PC (36:5) (16:0/20:5) | 17.64 | 0.11 | 42.51 | 3.06 |
| PC (38:4) (18:0/20:4) | 20.58 | 0.22 | 33.15 | 6.64 |
| PC (36:4) (16:0/20:4) | 22.62 | 0.18 | 59.68 | 3.60 |
| PC (38:5) (16:0/22:5) | 30.88 | 1.19 | 61.75 | 4.34 |
| PC (38:5) (18:1/20:4) | 30.88 | 1.19 | 61.75 | 4.34 |
| PC (30:0) (14:0/16:0) | 52.68 | 1.91 | 81.08 | 4.76 |
| PC (36:3) (18:1/18:2) | 48.52 | 1.73 | 125.49 | 8.98 |
| PC (36:1) (18:0/18:1) | 80.15 | 1.55 | 153.84 | 8.88 |
| PC (32:0) (16:0/16:0) | 183.71 | 3.77 | 232.08 | 9.17 |
| PC (34:2) (16:0/18:2) | 126.61 | 4.40 | 289.96 | 18.86 |
| PC (34:2) (16:1/18:1) | 126.61 | 4.40 | 289.96 | 18.86 |
| PC (36:2) (18:1/18:1) | 138.85 | 4.87 | 307.13 | 20.53 |
| PC (32:1) (14:0/18:1) | 172.51 | 4.09 | 375.94 | 21.72 |
| PC (32:1) (16:0/16:1) | 172.51 | 4.09 | 375.94 | 21.72 |
| PC (34:1) (16:0/18:1) | 335.55 | 10.51 | 661.80 | 38.41 |

| | | | | |
|---|---|---|---|---|
| PC (26:0) | 0.78 | 0.20 | 1.03 | 0.06 |
| PC (37:2) | 1.63 | 0.13 | 3.26 | 0.18 |
| PC (37:5) | 1.78 | 1.28 | 4.32 | 0.34 |
| PC (33:0) | 3.04 | 0.08 | 3.76 | 0.23 |
| PC (38:1) | 3.26 | 0.21 | 4.51 | 0.30 |
| PC (31:0) | 4.85 | 0.55 | 7.07 | 0.31 |
| PC (35:1) | 5.51 | 0.17 | 10.02 | 0.73 |
| PC (31:1) | 4.69 | 3.44 | 11.59 | 0.81 |
| PC (35:2) | 5.36 | 0.74 | 11.96 | 0.68 |
| PC (33:2) | 8.26 | 0.98 | 15.02 | 1.08 |
| PC (38:2) | 9.03 | 0.78 | 17.67 | 0.96 |
| PC (40:5) | 9.23 | 1.33 | 18.83 | 1.07 |
| PC (30:1) | 4.47 | 3.34 | 22.54 | 1.27 |
| PC (34:0) | 22.72 | 1.81 | 28.13 | 1.23 |
| PC (34:3) | 25.14 | 1.75 | 30.07 | 1.16 |
| PC (34:4) | 23.37 | 1.18 | 54.23 | 3.07 |
| PC (38:3) | 19.72 | 0.27 | 58.13 | 4.94 |
| PC (32:2) | 19.05 | 4.72 | 70.11 | 3.62 |
| PC (30:0) | 54.16 | 5.64 | 89.42 | 4.79 |
| PC (36:3) | 40.99 | 17.73 | 137.04 | 8.69 |
| PC (36:1) | 98.11 | 8.07 | 174.02 | 9.88 |
| PC (32:0) | 196.47 | 1.29 | 257.32 | 12.33 |

| | | | | |
|---|---|---|---|---|
| PC (36:2) | 157.56 | 1.10 | 350.23 | 23.14 |
| PC (34:2) | 144.75 | 10.56 | 367.25 | 24.38 |
| PC (32:1) | 206.97 | 19.26 | 476.49 | 27.23 |
| PC (34:1) | 357.49 | 6.56 | 702.03 | 42.05 |
| PE (36:5) | 8.66 | 3.94 | 8.81 | 0.94 |
| PE (38:6) | 9.48 | 2.49 | 20.67 | 1.70 |
| PE (40:5) | 10.19 | 7.28 | 24.74 | 2.65 |
| PE (38:4) | 18.64 | 1.00 | 41.43 | 5.40 |
| PE (40:7) | 13.82 | 3.22 | 27.99 | 2.58 |
| PE (38:5) | 26.37 | 3.40 | 44.77 | 4.71 |
| PE (34:1) (16:0/18:1) | 27.01 | 20.02 | 66.33 | 6.74 |
| PE (36:1) (18:0/18:1) | 70.79 | 23.90 | 76.87 | 7.31 |
| PE (36:2) (18:1/18:1) | 46.17 | 7.45 | 86.00 | 8.97 |
| PI (36:4) | 12.89 | 0.32 | 22.13 | 0.74 |
| PI (36:4) (16:0/20:4) | 12.89 | 0.32 | 22.13 | 0.74 |
| PI (36:1) (18:0/18:1) | 14.31 | 0.68 | 29.08 | 1.47 |
| PI (36:2) | 32.32 | 1.73 | 63.09 | 2.78 |
| PI (36:2) (18:1/18:1) | 32.32 | 1.73 | 63.09 | 2.78 |
| PI (38:5) | 73.34 | 3.68 | 101.82 | 4.30 |
| PI (38:3) (18:0/20:3) | 77.86 | 4.01 | 132.62 | 5.61 |
| PI (38:4) (18:0/20:4) | 163.50 | 8.06 | 275.08 | 13.33 |
| PS (34:1) (16:0/18:1) | 151.81 | 19.38 | 105.07 | 3.48 |
| PS (36:2) (18:0/18:2) | 155.61 | 18.60 | 105.06 | 3.12 |
| PS (36:2) (18:1/18:1) | 155.61 | 18.60 | 105.06 | 3.12 |
| PS (40:5) (18:0/22:5) | 172.76 | 19.78 | 129.69 | 3.79 |
| PS (36:1) (18:0/18:1) | 708.73 | 78.53 | 462.19 | 19.44 |
| SM (d35:1) | 2.42 | 0.63 | 2.32 | 0.07 |
| SM (d40:2) | 9.64 | 1.96 | 6.67 | 0.20 |
| SM (d32:1) | 10.47 | 2.45 | 8.54 | 0.51 |
| SM (d36:1) | 11.18 | 1.43 | 8.45 | 0.75 |
| SM (d42:1) | 21.53 | 7.48 | 11.98 | 0.92 |
| SM (d34:0) | 19.67 | 1.56 | 16.00 | 0.71 |
| SM (d42:3) | 37.67 | 9.15 | 27.07 | 0.96 |
| SM (d34:2) | 32.24 | 5.61 | 32.65 | 1.07 |

| | | | | |
|---|---|---|---|---|
| SM (d42:2) | 104.34 | 28.61 | 65.10 | 3.15 |
| SM (d34:1) | 438.87 | 19.18 | 323.70 | 7.71 |
| TG (56:7) | 1.54 | 0.09 | 3.50 | 0.24 |
| TG (56:7) (16:0/18:1/22:6) | 1.54 | 0.09 | 3.50 | 0.24 |

**Table S3.** Osmolarities of CDV hosting solutions. Measured data are mean ± standard deviations from 3 individual measurements.

| Hosting solution | MEAN [mOsm/kg] | STD |
|---|---|---|
| RPMI 1640 without supplements | 282.67 | 11.56 |
| PRMI 1640 without supplements + 10 μM cytochalasin B | 296.67 | 2.49 |
| PBS pH 7.4 (manufacturer's data) | 297.50 | 17.5 |
